# ATAC-seq and MNase-seq Detect Distinct Modes of Chromatin Accessibility

**DOI:** 10.64898/2026.09.07.749956

**Authors:** Shane Stoeber, Mitchell Godin, Lu Bai

**Affiliations:** Department of Biochemistry and Molecular Biology, The Pennsylvania State University, University Park, PA 16802, USA; Center for Eukaryotic Gene Regulation, The Pennsylvania State University, University Park, PA 16802, USA; Department of Physics, The Pennsylvania State University, University Park, PA 16802, USA

## Abstract

Chromatin accessibility shapes the ability of transcription factors (TFs) and the transcriptional machinery to engage genomic DNA and therefore plays a central role in gene regulation. Two widely used approaches for profiling chromatin accessibility are micrococcal nuclease (MNase)-seq and assay for transposase-accessible chromatin (ATAC)-seq. ATAC-seq peaks are often thought to be equivalent to nucleosome-depleted regions (NDRs) that are defined by MNase-seq; however, these two measurements have not been systematically compared. Here, we perform a side-by-side comparison of ATAC-seq and MNase-seq in budding yeast and find a large discrepancy between ATAC-seq peaks and MNase-defined NDRs. We show that this discrepancy is not primarily due to the intrinsic differences between MNase and Tn5 enzymatic activity. Instead, ATAC-seq peaks and NDRs capture distinct chromatin states. Specifically, ATAC-seq peaks are enriched at dynamic nucleosomes associated with transcriptional co-regulators, including SAGA and SWI/SNF, whereas ATAC− NDRs mark more static nucleosome-free regions at promoters. Depletion of SWI/SNF reduces ATAC-seq signals without affecting most NDRs. Generation of NDRs and ATAC-seq peaks requires distinct TF properties, and native TFs differ in their ability to produce these two types of open chromatin. Finally, we find that the functional distinctions between ATAC-seq peaks and NDRs are widespread across eukaryotic species, including human cells. Together, our results provide new insights into the biological meaning of chromatin accessibility measured by these two assays.

## Introduction

Chromatin accessibility plays an essential role in gene regulation by modulating the capability of transcription factors (TFs) and transcription machinery to engage genomic DNA. Mapping nucleosome occupancy and chromatin accessibility is therefore critical in understanding chromatin organization and transcriptional control. Two widely used experimental approaches to profile genome-wide chromatin accessibility are MNase-seq and ATAC-seq^1^. MNase-seq employs Micrococcal Nuclease (MNase), an endo-exonuclease that preferentially digests linker DNA relative to nucleosomal DNA. MNase-seq is typically performed under conditions in which most chromatin is fragmented into mono-nucleosome-sized particles^2^, and sequencing these fragments provides a high-resolution map of nucleosome positioning across the genome. Long gaps between nucleosomes, typically between 80 to 300 bp, are annotated as nucleosome-depleted regions (NDRs). In contrast, ATAC-seq utilizes the Tn5 transposase to simultaneously cleave accessible DNA and ligate adaptor sequences in a process known as tagmentation^3^. Amplifying and sequencing tagmented DNA fragments generates ATAC-seq peaks that correspond to chromatin regions with high Tn5 accessibility.

While MNase-seq provides higher resolution nucleosome positioning information, it requires a large number of cells and high sequencing depth to achieve robust signals, making it challenging to apply to species with large genomes, including human and mouse^4^. Most MNase-seq, therefore, are performed in model organisms with smaller genome sizes, like budding yeast. ATAC-seq is complimentary in this regard, providing information on accessible regions without sufficient resolution to annotate nucleosome positioning, but requires much less cells with lower sequencing coverage^5^. ATAC-seq has therefore become the predominant chromatin accessibility assay for mammalian cells. For example, the ENCODE database contains ∼800 bulk and single-nuclei ATAC-seq datasets, but only two MNase-seq datasets.

Because these two methods have largely been applied to different samples, they have not been directly compared side-by-side under matched experimental conditions to our knowledge. It is often assumed that NDRs detected by MNase-seq are equivalent to ATAC-seq peaks, as both are indicative of open chromatin^6,7^. Consequently, increases in ATAC-seq signals are frequently interpreted as nucleosome remodeling or eviction. These assumptions have not been rigorously tested. Here, we address this gap by performing a direct comparison of MNase-seq and ATAC-seq in budding yeast. Besides its small genome size, yeast nucleosomes tend to be regularly-phased and well-positioned, allowing us to rigorously annotate genome-wide NDRs^8–10^. In contrast, nucleosome positioning in higher eukaryotes is much fuzzier^11^. The chromatin organization in yeast therefore provides a system in which MNase-defined NDRs and ATAC-seq peaks can be compared with reduced ambiguity.

This comparison reveals a substantial discrepancy between the two assays. Only ∼20% of NDRs have associated ATAC-seq peaks, and only ∼half of ATAC-seq peaks overlap with NDRs. We found that this discrepancy is not primarily due to intrinsic differences between MNase and Tn5 but reflects chromatin properties that render differential susceptibility to the two enzymes. ATAC-seq peaks are enriched at dynamic nucleosomes associated with transcriptional co-regulators, including SAGA and SWI/SNF, whereas NDRs without ATAC signal mark more static nucleosome-free regions at promoters. Depletion of SWI/SNF, but not RSC or the SAGA complex, reduces genome-wide ATAC-seq signals. Generation of NDRs and ATAC-seq peaks requires distinct TF properties, and native yeast transcription factors (TFs) differ in their ability to produce these two types of open chromatin. Finally, we show that the discrepancy between ATAC-seq peaks and NDRs, as well as their differential association with transcriptional cofactors, is widespread across eukaryotic species, including human cells. Together, these findings challenge the common interpretation of ATAC-seq peaks as simple proxies for nucleosome depletion and instead suggest that ATAC-seq and MNase-seq reveal distinct modes of chromatin accessibility.

## Results

### ATAC-seq peaks and NDRs mapped by MNase-seq show striking discrepancies

We performed ATAC-seq and MNase-seq measurements using matched yeast cultures (all strains are listed in **Table S1**). Standard methods were used to analyze these data and identify ATAC-seq peaks and NDRs (**Methods**). Since nucleosome depletion over yeast terminators tend to be weak^12^, we excluded NDRs located in convergent intergenic regions or >500 bp from a transcription start sites (TSSs). After this filtering, we identified 1242 ATAC peaks and 3504 NDRs, which are clustered into three groups: 1) “ATAC+ NDRs” or “overlapped”, where ATAC-seq peaks coincide with NDRs (*N* = 672), 2) “NDR only”, NDRs with no significant ATAC peaks (*N* = 2832), and 3) “ATAC only”, ATAC peaks with no corresponding NDRs (*N* = 603) (**Figure 1A**). In summary, about half of ATAC peaks overlap with NDRs and conversely, only ∼ 20% of NDRs show significant ATAC signals. Importantly, our ATAC-seq and MNase-seq data, as well as the clustered pattern in **Figure 1A**, are highly consistent with previously published datasets^13,14^ (**Figure S1A & B**). The discrepancy of the two assays is therefore not a technical artifact of our measurements but represents a reproducible feature of the two assays.

**Figure 1.**
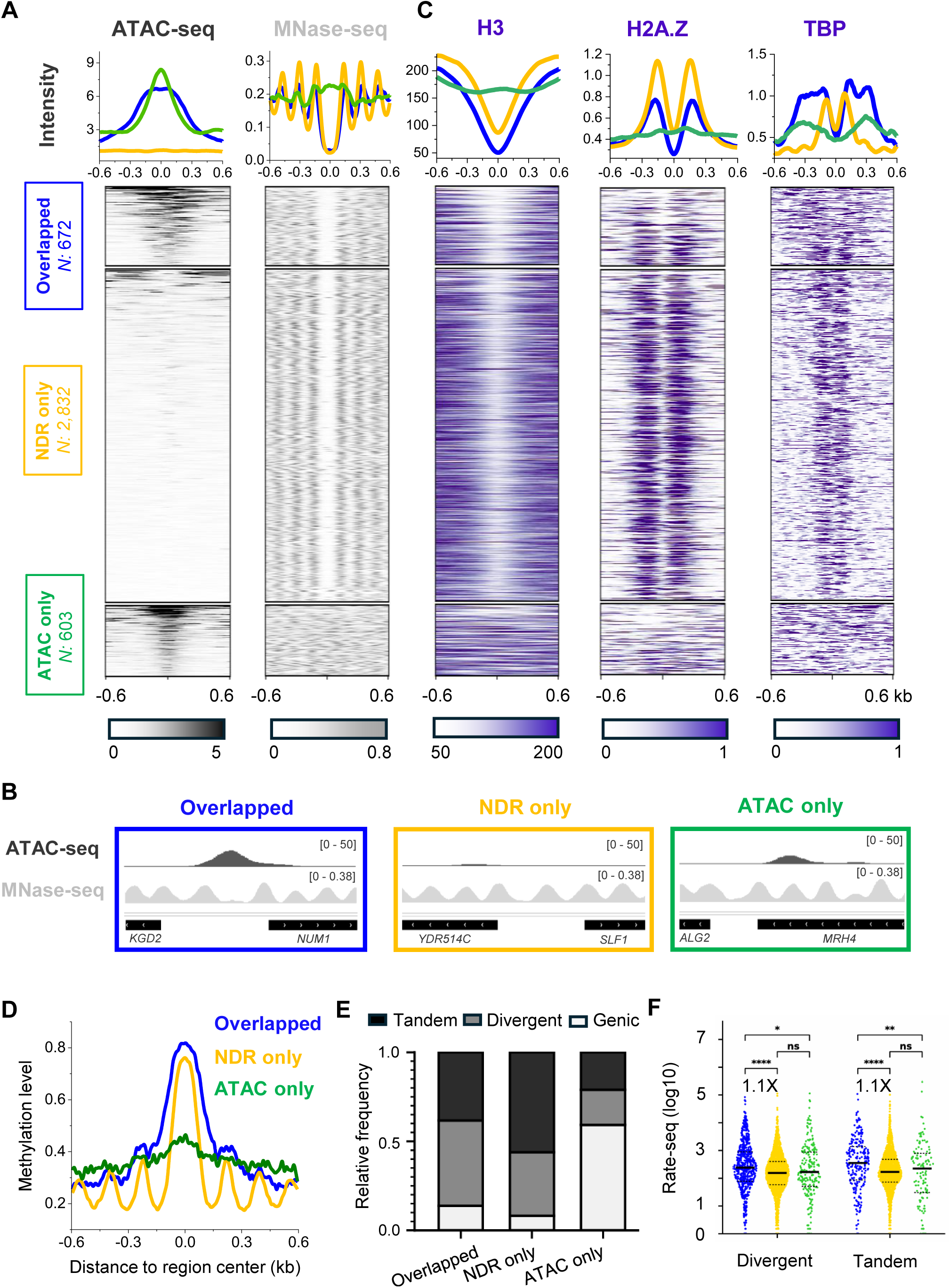
Discrepancy between ATAC-seq peaks and NDRs. **A)** Comparison between our ATAC-seq and MNase-seq data in budding yeast. Regions are clustered into three groups: “overlapped”, in which ATAC-seq peaks coincide with NDRs; “NDR only”, in which NDRs lack significant ATAC-seq peaks; and “ATAC only”, in which ATAC-seq peaks lack corresponding NDRs. **B)** Representative ATAC-seq and MNase-seq tracks of the three groups shown in panel A. **C)** H3, H2A.Z, and TBP ChIP-seq data^17^ in the three clusters. Rows are ordered as in panel A. **D)** Accumulated DNA methylation level over the three clusters based on published Fiber-seq data^19^. **E)** Genomic location of the overlapped, NDR only, and ATAC only regions. Tandem and divergent: intergenic regions between tandem and divergent genes, Genic: over gene body. **F)** RATE-seq counts of nascent transcripts^20^ driven by the promoters associated with each group. For divergent promoters, both flanking genes are included; for tandem promoters, only the downstream gene is included. ****: P-value <0.00001, ns, not significant.

### NDRs exhibit features consistent with reduced nucleosome occupancy, regardless of ATAC signals

Given the discrepancy observed above, we next asked which group of open chromatin best reflects the underlying “ground truth” of nucleosome occupancy. Do regions with conflicting signals (NDR only and ATAC only) represent bona fide open regions with nucleosome depletion? To address this, we analyzed independent datasets over the three clustered regions that provide orthogonal measures of nucleosome occupancy. First, H3 ChIP-seq^15^ confirms that all NDRs, regardless of ATAC-seq signal, display strong depletion of histone H3. By contrast, ATAC only regions show no reduction in H3 signal (**Figure 1C**). Second, because the SWR1 chromatin remodeling complex targets NDRs to deposit H2A.Z into the flanking nucleosomes^16^, H2A.Z enrichment^17^ serves as an additional marker of NDRs. Both overlapped and NDR only regions show strong H2A.Z enrichment, whereas ATAC only regions are largely devoid of H2A.Z signal (**Figure 1C**). This pattern is further supported by TBP (SPT15) binding. As an essential transcription initiation factor, TBP binding is generally incompatible with nucleosome occupancy^18^. TBP ChIP-exo data^17^ show enrichment in the first two clusters but not in the ATAC only cluster (**Figure 1C**). Finally, DNA methylation measurements provide an independent readout of chromatin accessibility. Following treatment with exogenous DNA methyltransferase, accessible DNA is expected to acquire higher levels of methylation. Analysis of previously published Fiber-seq data^19^ reveals high DNA methylation levels in NDRs ± ATAC-seq signal, but substantially lower methylation in ATAC only regions (**Figure 1D**). Overall, these findings provide strong support for the NDRs detected by MNase-seq and indicate that ATAC-seq has variable sensitivity towards different NDRs.

We next examined the genomic distribution of the three clusters. Most overlapped and NDR only regions are located in promoters between divergent or tandem genes, while ∼60% of the ATAC only regions reside within gene bodies (**Figure 1E**). To assess transcriptional activity, we used published RATE-seq data^20^ to quantify nascent transcripts from tandem and divergent promoters associated with each ATAC/NDR group. Promoters in all three groups show a wide range of activities, with overlapped promoters having a slightly higher median transcript level (∼1.1X) than NDR only / ATAC only promoters (**Figure 1F**). A similar analysis using RNA-seq data^21^ reveals the same trend (**Figure S1C**). These results indicate that overlapped, NDR only, and ATAC only regions can all serve as functional promoters that can drive active transcription. In addition, ATAC-seq peaks can arise from nucleosome-covered or partially covered regions, many of which are located within gene bodies.

#### ATAC signals are not caused by “fuzzy” nucleosome positioning

Although most yeast nucleosomes are well-phased, a subset exhibits fuzzy positioning (**Figure S2A**). Our initial visual inspection showed that some ATAC-seq peaks overlap fuzzy nucleosome arrays, raising the possibility that the fuzziness reflects nucleosome dynamics that allow Tn5 to access these sites. To test this idea, we used the previously established DANPOS algorithm^22^ to calculate fuzziness scores for nucleosomes flanking ATAC+ and ATAC− NDRs, as well as nucleosomes near the centers of ATAC only regions (**Figure S2B**). This analysis shows that fuzziness scores are only modestly higher in regions with ATAC-seq peaks (**Figure S2C**). Moreover, many ATAC-seq peaks overlap well-positioned nucleosome arrays, whereas many fuzzy arrays in the genome lack detectable ATAC-seq signals (**Figure S2A**). We therefore conclude that fuzzy nucleosome positioning is neither necessary nor sufficient to generate ATAC-seq peaks.

### Discrepancy between ATAC-seq peaks and NDRs is not mainly caused by the intrinsic difference between MNase and Tn5

To understand the discrepancy between ATAC-seq peaks and MNase-defined NDRs, we considered the inherent sequence biases of the two enzymes. Previous studies have shown that MNase preferentially cleaves AT-rich DNA, whereas Tn5 prefers GC-rich sequences^23,24^. Consistent with these opposite sequence biases, base composition analysis shows that NDR only regions have higher AT content than the other two groups (**Figure 2A**). To further evaluate this effect, we scanned the sequences in the three groups using a previously reported Tn5 position weight matrix (**Figure 2B**)^25^. The Tn5 motif occurs at higher frequencies in the two ATAC+ groups, but the overall fraction of ATAC-seq peaks containing a strong Tn5 consensus motif remains small (**Figure 2B**).

**Figure 2.**
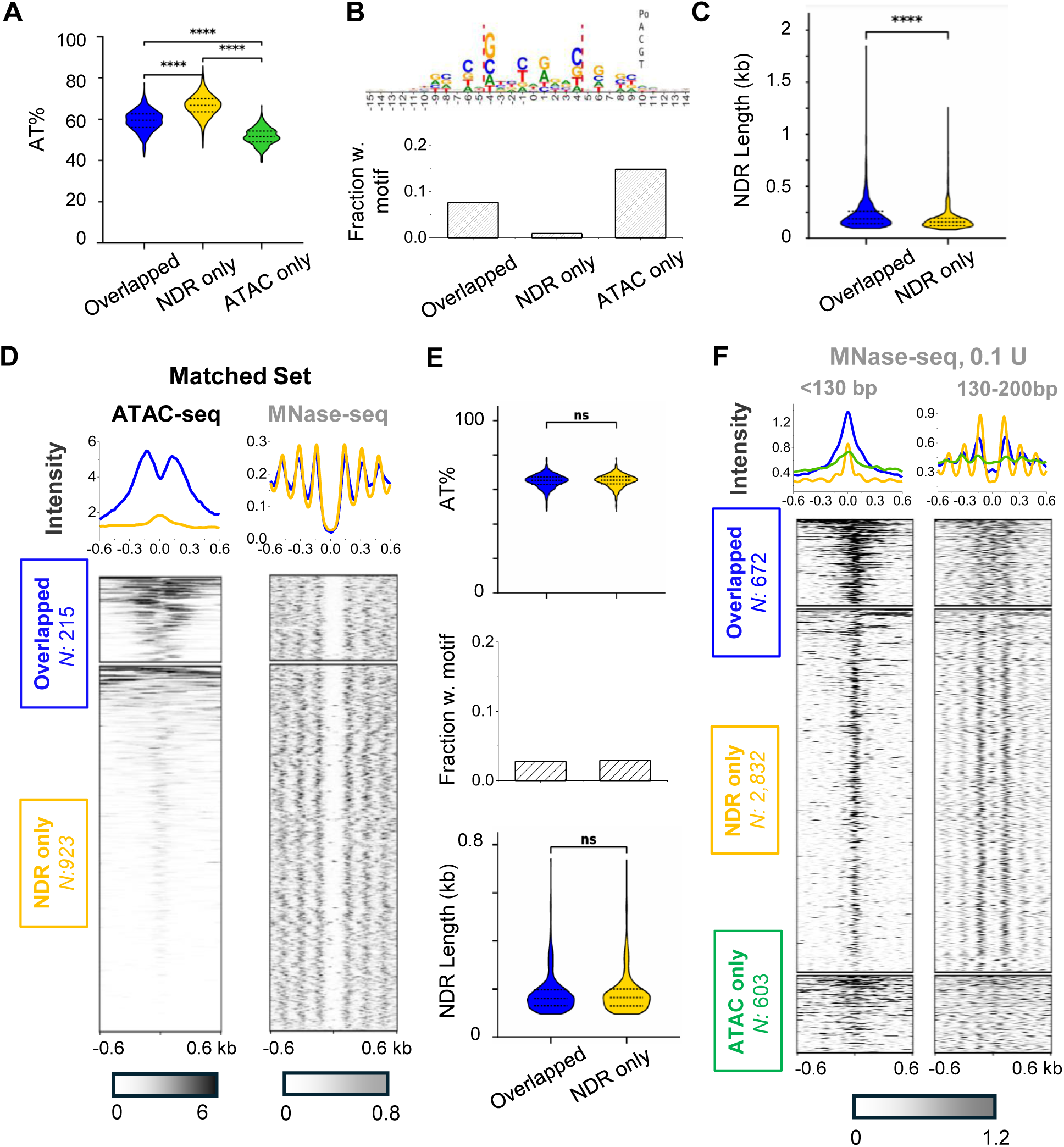
Sequence composition and NDR length do not fully explain the discrepancy between ATAC-seq peaks and MNase-defined NDRs. **A)** AT content (%) of regions in overlapped, NDR only, and ATAC only clusters. For overlapped and NDR only regions, AT content is calculated across each NDR. For ATAC only regions, AT content is calculated across each ATAC-seq peak. ****: P-value <0.00001 (same as below). **B)** Tn5 motif^53^ and its occurrence frequency in the three clusters. **C)** NDR length distribution in overlapped vs NDR only regions. **D & E)** Matched subset of overlapped and NDR only regions with statistically the same AT content, motif frequency, and NDR length distributions. ns, not significant. **F)** MNase-seq performed at low MNase concentration (0.1U) separated by fragment size. Left: sub-nucleosomal reads (<130bp), and right: nucleosomal reads (130-200 bp). The sub-nucleosomal MNase-seq signal closely resembles the ATAC-seq signal.

Another factor that may contribute to this discrepancy is the length of the NDR. Detection of ATAC-seq signal requires two tagmentation events separated by a sufficient distance so that the resulting DNA fragment can survive the size selection during library preparation. Therefore, ATAC-seq is inherently biased against short NDRs, which may fail to produce detectable fragments even if cleaved multiple times by Tn5. Consistent with this prediction, we compared the length of NDRs with or without ATAC signal and found the former to be moderately longer (median length 186bp vs 155bp) (**Figure 2C**). This difference is also visible in the MNase-seq metaprofiles, where the +1 and −1 nucleosomes are separated by a slightly wider gap across the overlapped regions (**Figure 1A**).

The overall differences in base composition and NDR length are modest. Moreover, H3 ChIP-seq, H2A.Z ChIP-exo, and fiber-seq measurements do not share the same biases, yet they agree well with MNase-seq results (**Figure 1A & C**). We therefore suspected that sequence bias and NDR size cannot fully account for the discrepancy between ATAC-seq peaks and NDRs. Consistent with this idea, we selected two “matched” NDR subgroups with nearly identical AT content, Tn5 motif incidence, and NDR length distributions, but with markedly different ATAC-seq signals (215 ATAC+ and 923 ATAC−) (**Figure 2D & E**). Thus, for a substantial fraction of NDRs, the presence or absence of ATAC-seq signal must be determined by additional features. We use this matched subset in subsequent analyses to control for potential technical confounders.

We reasoned that if the intrinsic differences between MNase and Tn5 contribute only modestly to the observed discrepancy, MNase-seq data should be more similar to ATAC-seq when performed at lower digestion frequencies. To test this, we performed a low-concentration MNase-seq (0.1U) and analyzed nucleosomal fragments (130–200 bp) and sub-nucleosomal fragments (<130 bp) separately (**Figure 2F, S3**). Under the standard high-MNase condition, these two fragment classes show similar distributions (**Figure S3C & D**), indicating that most sub-nucleosomal fragments arise from over-digested nucleosomes. In contrast, at low MNase concentration, the two fragment sizes generate highly contrasting heatmaps: nucleosomal fragments remain at their canonical locations, while sub-nucleosomal fragments are concentrated near the center of NDRs and ATAC only regions, closely resembling ATAC-seq patterns (**Figure 2F**). Such similarity supports the notion that the differences observed between the two assays are driven less by intrinsic differences between Tn5 and MNase, and more by the digestion regime and fragment size selection. These results also suggest that overlapped, NDR only, and ATAC only regions have unique properties that render different sensitivity to both MNase and Tn5.

### ATAC-seq peaks arise from tagmentation over broad regions, likely inside dynamic nucleosomes

Two intriguing observations can be made from **Figure 2F**. First, the +1/−1 nucleosomes flanking ATAC− NDRs are equally, if not more, detectable than the ones flanking ATAC+ NDRs. This suggests that these two groups of NDRs, despite their different ATAC profiles, are both susceptible to MNase digestion even at a low cutting frequency. Second, although ATAC only regions are largely nucleosomal, they can yield sub-nucleosomal fragments when MNase is applied at a low concentration.

To reconcile these patterns, we analyzed the positions of Tn5 tagmentation events (**Methods**). In contrast to whole-fragment pile-up, fragment edges (tagmentation sites) provide a more direct readout of local Tn5 accessibility. **Figure 3A** shows the tagmentation sites over two NDRs of similar size, one with strong ATAC-seq signal and one with minimal signal. It turns out that the latter can be tagmented, but all the tagmentation events are confined to a narrow window within the NDR. In contrast, for the ATAC+ NDR, tagmentation is spread across a broader region inside the flanking nucleosomes. This difference is also evident in averaged profiles of tagmentation frequency, where it spans a wider range in the overlapped regions (**Figure 3B**). The central ∼2 nucleosomes in ATAC only regions also show strong tagmentation signals (**Figure 3B**). These distinct tagmentation patterns can explain the difference in the cumulative ATAC-seq signals. For NDR only sites, even when local tagmentations occur frequently, the narrow accessible window generates mostly short fragments that are likely lost during library preparation due to size selection (**Figure 3C**). By contrast, broader tagmentation at overlapped regions increases the probability of generating longer fragments that are retained and sequenced.

**Figure 3.**
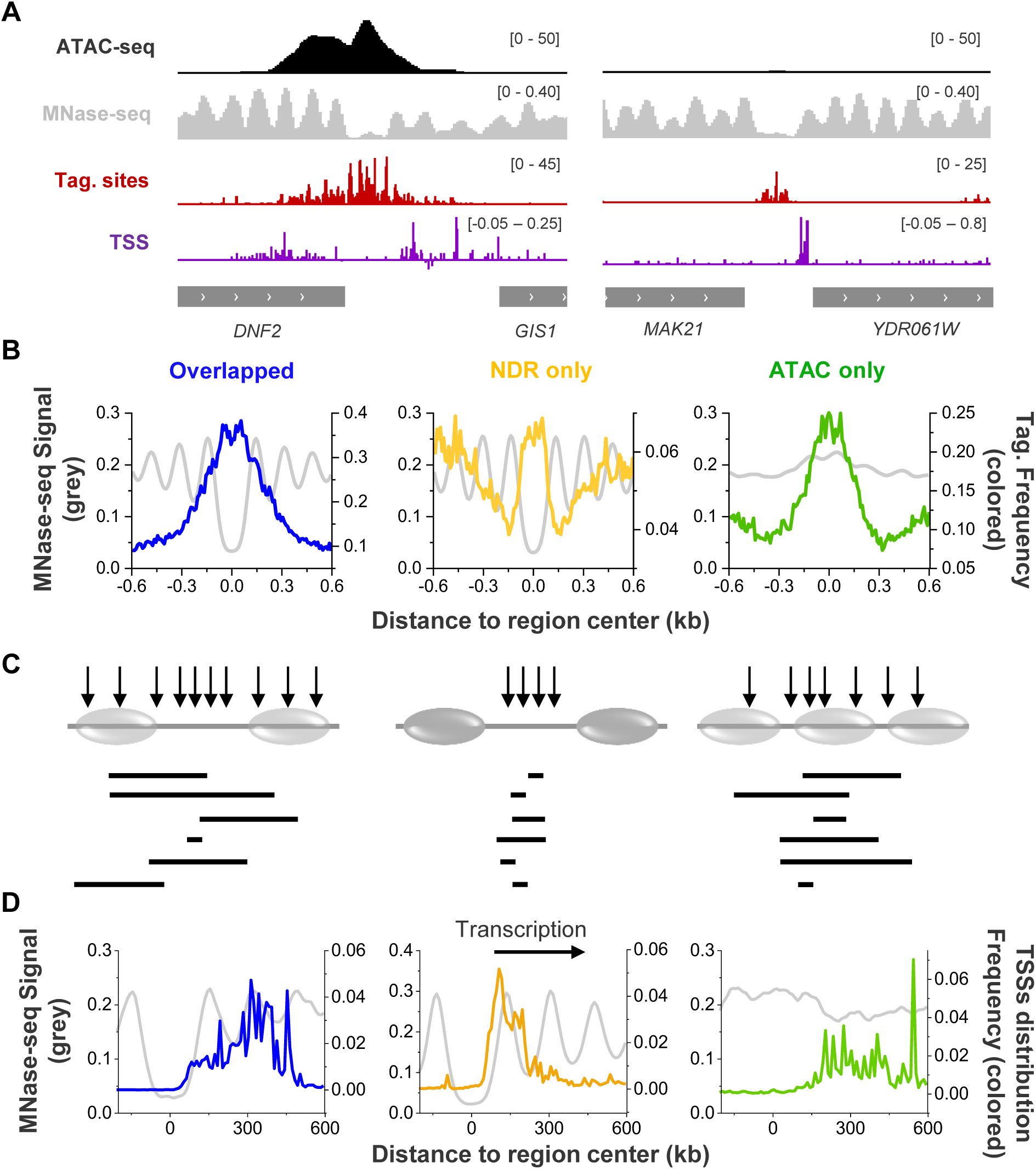
ATAC-seq peaks arise from broad tagmentation domains that often extend into flanking nucleosomes. **A)** Representative examples of ATAC-seq, MNase-seq, Tn5 tagmentation sites, and CAGE-seq (TSS)^28^ signals over one overlapped region (left) and one NDR only region (right). At the overlapped region, tagmentation extends beyond the NDR into flanking regions that also show nucleosome occupancy. **B)** Average nucleosome coverage (grey) and Tn5 tagmentation frequency (colored) across all overlapped, NDR only, and ATAC only regions. **C)** Cartoon illustrating how the tagmentation-permissive domain size can produce large differences in ATAC-seq signal. Solid / semi-transparent ovals represent static / dynamic nucleosomes. **D)** Distribution of TSSs for promoters in the three groups. All genes are reoriented towards right.

If ATAC-seq peaks largely reflect dynamic and accessible nucleosomes, how do these nucleosomes permit Tn5 tagmentation while remaining largely protected from MNase digestion? We suspect that this apparent conflict reflects population heterogeneity that is obscured by bulk averaging. One possible scenario is that, in a subset of cells, these nucleosomes may be destabilized or even evicted, creating transiently expanded NDRs that contribute disproportionately to the ATAC-seq signal. In other cells, the same nucleosomes remain intact, producing apparent protection in the MNase-seq assay.

To test the dynamic nucleosome model, we analyzed transcription start site (TSS) distributions at ATAC+ and ATAC− promoters. In yeast, Pol II selects TSSs through a scanning process, in which Pol II catalytic activity kinetically competes with translocation to determine the probability of initiation at each position^26^. TSSs often reside within the nucleosome immediately downstream of the promoter NDR^27^, presumably because this nucleosome slows Pol II movement and increases the probability of catalysis. We therefore reasoned that if ATAC+ promoters contain dynamic nucleosomes, Pol II should be able to scan further into nucleosome-occupied regions, resulting in broader and more downstream-shifted TSS distributions. Indeed, for the examples shown in **Figure 3A**, TSSs near *GIS1* penetrate more into the nucleosomes downstream of NDR, whereas TSSs near *YDR061W* are confined at the edge of the NDR. To determine if this trend is general, we plotted the TSS profiles for overlapped, NDR only, and ATAC only promoters based on previously published CAGE-seq data^28^ (**Methods**). Consistent with the hypothesis above, overlapped promoters show higher TSS usage within nucleosomes +2 or +3, while NDR only promoters initiate primarily within the +1 nucleosome (**Figure 3D & S3E**). ATAC only promoters also show broadly distributed TSSs downstream of the ATAC-seq peak center. Together, these data support the idea that nucleosomes in overlapped and ATAC only regions tend to be dynamic, and ATAC signals are particularly sensitive to these dynamic nucleosomes.

### ATAC-seq peaks strongly correlate with SAGA and SWI/SNF binding

We next asked which factors may drive nucleosome dynamics that give rise to ATAC-seq signals. Taking advantage of previously published ChIP-seq and ChEC-seq datasets^17,29^, we first evaluated the differential enrichment of transcription machinery, sequence-specific TFs, and regulatory co-factors in overlapped, NDR only, and ATAC only regions. Surprisingly, unlike TBP and H2AZ, which bind inside or close to NDRs (**Figure 1C**), many TFs and co-factors, including SAGA and SWI/SNF, show binding patterns highly correlated with ATAC-seq signals, including the ATAC only regions (**Figure 4A & S4A**).

**Figure 4.**
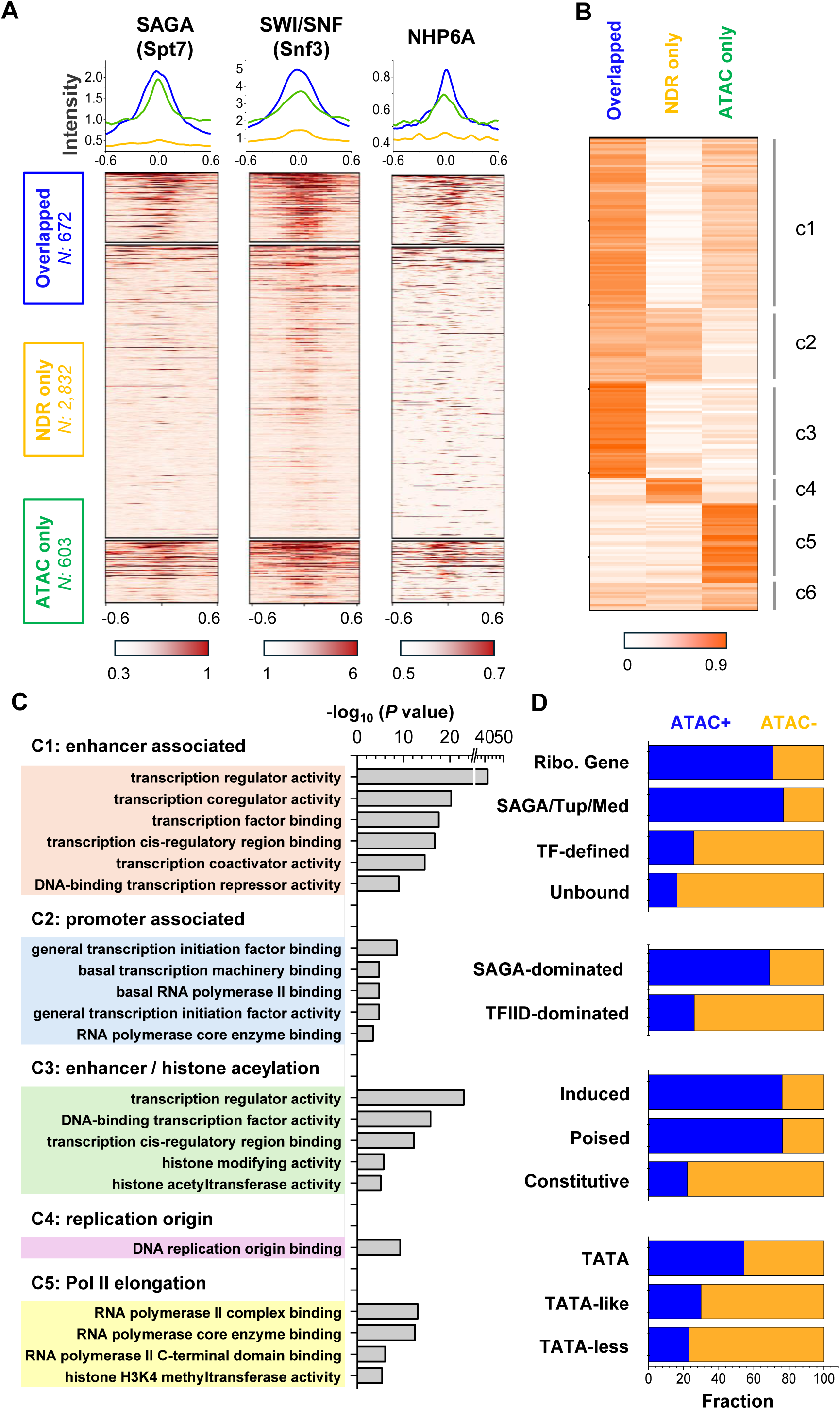
Factors preferentially bind to ATAC-seq peaks vs NDRs show functional divergence. **A)** Spt7 (SAGA subunit)^17^, Snf3 (SWI/SNF subunit)^29^, and NHP6A binding data^17^ in overlapped, NDR only, and ATAC only regions. All three factors are preferentially enriched at ATAC-seq peaks, including both overlapped and ATAC only regions. **B)** K-means clustering of binding probabilities for 233 chromatin-associated factors over overlapped, NDR only, and ATAC only regions (**Table S2**). Clusters c1–c5 represent factors preferentially enriched at ATAC-seq peaks, NDRs, overlapped regions, NDR only regions, and ATAC only regions, respectively. Factors in c6 show little preference among the three region classes. **C)** Gene Ontology analysis of factors within each cluster shown in panel B. **D)** Association between ATAC+ or ATAC− promoters and previously annotated promoter features.

Motivated by this observation, we performed k-means clustering of binding profiles for > 200 factors (mostly based on previously published ChIP-exo data^17^) and identified five major binding patterns (**Figure 4B & S4A; Table S2**). Groups c1 and c2 capture factors that preferentially bind ATAC peaks (e.g. SAGA) or NDRs (e.g. TBP), respectively. Clusters c3–c5 correspond to factors selectively enriched in overlapped, NDR only, and ATAC only regions, and the rest are grouped into c6. GO analysis of factors in c1-c5 reveals different functional categories. Most notably, c1 is enriched for factors associated with gene regulation / enhancer-like activities, c2 with transcription initiation / promoter functions, c4 with replication origin-related processes, and c5 with activities over genes (**Figure 4C**). These findings indicates that promoters with or without ATAC signals bind to different factors and likely have functional divergence.

Yeast promoters have been classified into several categories based on TF and cofactor binding patterns. These include promoters that drive ribosomal genes (RG), promoters associated with cofactors SAGA/Tup/Mediator (STM), promoters bound by sequence-specific TFs but not by STM (TF-organized), and promoters with no detectable sequence-specific TF binding (unbound)^17^. Promoters have also been divided into inducible and constitutive groups based on transcription and recruitment of transcription machinery^30^. Inducible promoters tend to be SAGA-associated, TATA-containing, and enriched for sequence-specific TF binding, whereas constitutive promoters are typically TFIID-associated, TATA-less, and depleted of sequence-specific TF binding. To examine how these classifications relate to ATAC-seq signal, we divided previously classified promoter NDRs into ATAC+ and ATAC− groups (**Methods**). RG and STM promoters are strongly associated with ATAC-seq peaks, whereas TF-organized and unbound promoters are depleted of ATAC-seq signal (**Figure 4D**). The depletion of ATAC-seq peaks at TF-organized promoters is particularly informative, as it indicates that recruitment of some cofactors, rather than the binding of TFs, is critical for generating ATAC signal. Consistent with the enrichment of SAGA binding, transcription from ATAC+ promoters is also more sensitive to SAGA depletion (SAGA-dominated) (**Figure 4D**). Importantly, although SAGA binding strongly correlates with ATAC-seq signal, transcription does not: “induced” and “poised” genes, with the latter showing little Pol II and general transcription factor (GTF) recruitment, are both highly enriched for ATAC-seq peaks (**Figure 4D**). This is consistent with our observation in **Figure 1F** that promoters in all categories drive similar level of transcription on average. Finally, promoters containing a canonical TATA box tend to overlap ATAC-seq peaks, although the fold enrichment is lower than that observed for categories defined by SAGA binding.

We reasoned that the differential enrichment in ATAC vs NDR (c1 vs c2) for some factors could, at least in part, reflect sequence bias (**Figure 2B**). For example, TBP recognizes TATA box and thus would be expected to enrich in A/T-rich NDRs. To eliminate such bias, we repeated the binding enrichment analysis using the subset defined in **Figure 2D**, where both the NDR length and AT content are matched between ATAC ± NDRs. Under these matched conditions, most factors still bind more frequently in ATAC+ than ATAC− NDRs, with 36 factors showing statistically significant enrichment (fold change >2 and effective *P-*value < 0.01) (**Figure 5A**). SAGA and SWI/SNF binding are the most correlated with ATAC-seq signals, with seven SAGA subunits and four SWI/SNF subunits highly enriched in ATAC+ NDRs. This raises the possibility that SAGA and SWI/SNF promote ATAC peaks by generating unstable and dynamic nucleosomes.

**Figure 5.**
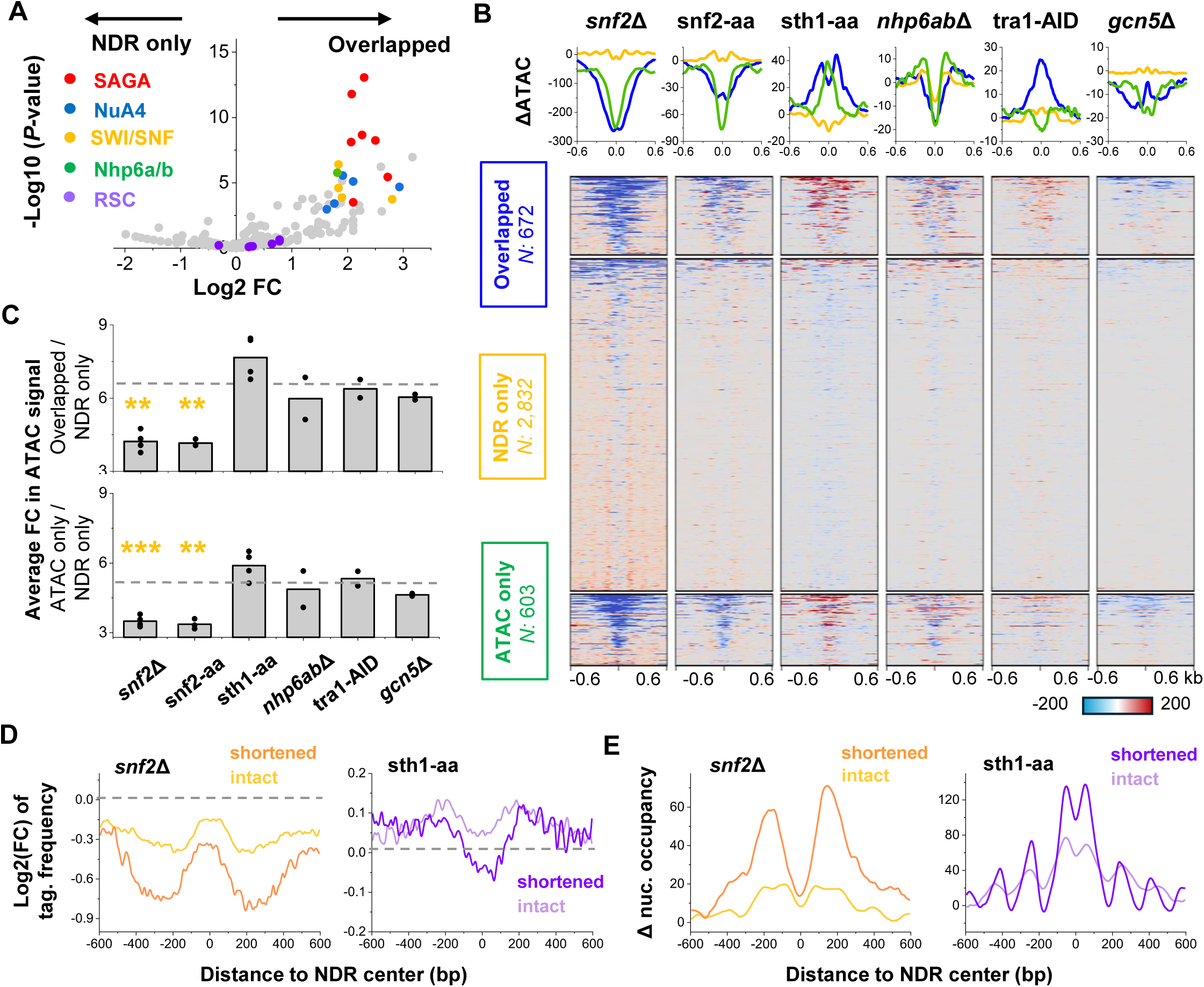
Depletion of SWI/SNF, but not RSC, significantly reduces ATAC-seq signals. **A)** Volcano plot showing differential enrichment of chromatin-associated factors at overlapped vs NDR only regions. The regions used here are AT-content- and length-matched, as defined in Figure 2D. Factors selected for perturbation experiments are highlighted with colored dots. **B)** Changes in ATAC-seq signal following genetic deletion, mutation, or rapid depletion of selected factors. “Δ” indicates genetic mutation or deletion; aa, anchor-away; AID, auxin-induced degradation. **C)** Average ATAC-seq signal at overlapped and ATAC only regions in each perturbation sample, normalized to the signal at NDR only regions. The dotted line indicates the WT level. **: P-value <0.01, ***: P-value <0.001. **D)** Average change in Tn5 tagmentation frequency near NDRs after *SNF2* mutation or Sth1 anchor-away. NDRs are analyzed separately based on whether they shrink or remain unchanged after remodeler disruption. **E)** Same analysis as in panel D, but for nucleosome occupancy measured by MNase-seq.

### Depletion of SWI/SNF, but not RSC, significantly reduces ATAC-seq signals

To test the causal relationship between factor binding and ATAC-seq profiles, we performed a series of deletion or rapid-depletion experiments to measure the resulting changes in ATAC-seq signals. Besides the SAGA and SWI/SNF complexes mentioned above, we disrupted three other chromatin-associated factors: 1) NuA4, another histone acetyltransferase complex, 2) Nhp6a/b, a pair of redundant high-mobility group proteins, and 3) RSC, an essential SWI/SNF-family remodeler. NuA4 and Nhp6a/b were selected because of their enrichment at ATAC+ NDRs (**Figure 5A**) and potential roles in nucleosome instability^31,32^. Although RSC subunits are not enriched at ATAC+ NDRs (**Figure 5A**), we included it because of its broad effect on genome-wide nucleosome positioning^33,34^. More specifically, we perturbed SAGA in two ways: by deleting *GCN5*, which encodes the catalytic subunit of the SAGA HAT module, or through auxin-induced degradation of Tra1, an essential SAGA subunit shown to interact with transcriptional activators^35^ (**Figure S5A**). Because Tra1 is a component in both SAGA and NuA4^36^, its depletion is expected to disrupt both complexes. SWI/SNF is also disrupted with two methods, either by introducing the E834K mutation into its catalytic subunit *SNF2*, which impairs its remodeling activity^37^, or by rapidly depleting Snf2 through anchor-away (**Figure S5B**). To examine the contribution of Nhp6 proteins, we utilized a *nhp6a*Δ *nhp6b*Δ double-deletion strain, given their redundant functions. Finally, we disrupted RSC by anchoring away its catalytic subunit, Sth1 (**Figure S5B**).

We quantified ATAC-seq signal changes in each perturbation relative to WT and presented the difference per region in **Figure 5B** and the average fold change in **Figure 5C** (**Methods**). Among the factors tested, SWI/SNF disruption produces the strongest effect: both the *SNF2* mutation and acute Snf2 depletion cause genome-wide reduction in ATAC-seq signal. This reduction is observed for both sub-nucleosomal fragments (<130 bp) and nucleosomal fragments (130–200 bp) (**Figure S5C**). In contrast, RSC depletion mildly increases ATAC-seq signals, with a pattern that is opposite to Snf2-aa (**Figure 5B & C**). The remaining disruptions of Gcn5, Tra1, and Nhp6a/b generate even weaker effects, although Tra1 degradation causes a small increase in ATAC signal, and Nhp6a/b deletion slightly reduces the sub-nucleosomal fragments and increases the nucleosomal fragments (**Figure 5B & S5C**).

To further understand these observed effects, we evaluated the changes in Tn5 tagmentation frequency upon SWI/SNF or RSC depletion. We focused our analysis on overlapped regions because 1) ATAC-seq signals over these regions show large responses to remodeler depletion, and 2) they can be separated into “shortened NDRs”, which shrink upon remodeler depletion, and “intact NDRs”, whose sizes remain largely the same. SWI/SNF depletion lowers tagmentation frequency in both groups, with a stronger effect towards the former (**Figure 5D**). Importantly, the largest decrease occurs over the flanking nucleosomes rather than within the NDRs themselves. This pattern closely matches nucleosome occupancy changes measured by MNase-seq^29^, which shows that SWI/SNF depletion leads to global increases in the occupancy of NDR-flanking nucleosomes without filling in the NDRs (**Figure 5E, S5D**). Overall, these data indicate that SWI/SNF primarily promotes Tn5 accessibility by destabilizing NDR-flanking nucleosomes.

In comparison, RSC depletion produces a smaller effect in the opposite direction. At NDRs that shorten upon RSC depletion, tagmentation decreases slightly near the center of NDRs (**Figure 5D**). This effect is surprisingly small given the strong increase in nucleosome occupancy within these NDRs (**Figure 5E, S5D**). Near both shortened and intact NDRs, tagmentation frequency shows a minor increase in the flanking nucleosomes (**Figure 5D**), indicating that RSC reduces the accessibility of these nucleosomes. This could be achieved by RSC covering up these regions sterically or by competing with SWI/SNF for binding at the same sites.

### Transcription factors have differential ability to generate NDR vs ATAC signals

The discrepancy between NDRs and ATAC-seq peaks is also important when considered in the context of TFs. In particular, a subset of sequence-specific TFs named as pioneer factors (PF) can bind nucleosomes and create accessible chromatin^38,39^. However, “accessible chromatin” is defined in different ways in the literature, either as MNase-defined NDRs or as ATAC-seq peaks. Based on our findings above, we suspect that TFs may differ in their ability to generate NDRs versus ATAC-seq signals, and PFs defined by these two measurements are not necessarily equivalent.

To test this idea, we first used an artificial system in which a bacterial TF LexA is expressed in budding yeast and targeted to a well-positioned nucleosome at the *HO* promoter (*HOpr*) through two engineered LexA recognition motifs (**Figure 6A**). Consistent with our previous study^40^, LexA can invade into this nucleosome and generate a 120–150 bp NDR (**Figure 6A**), comparable to the average size of native NDRs. We then asked whether this LexA-induced NDR also produces an ATAC-seq signal by carrying out Tn5 tagmentation followed by stacking qPCR near the LexA sites (**Methods**). Despite near-complete depletion of the central nucleosome, little ATAC signal is detected across this region (**Figure 6A**). Thus, LexA-induced open chromatin falls into the NDR only category.

**Figure 6.**
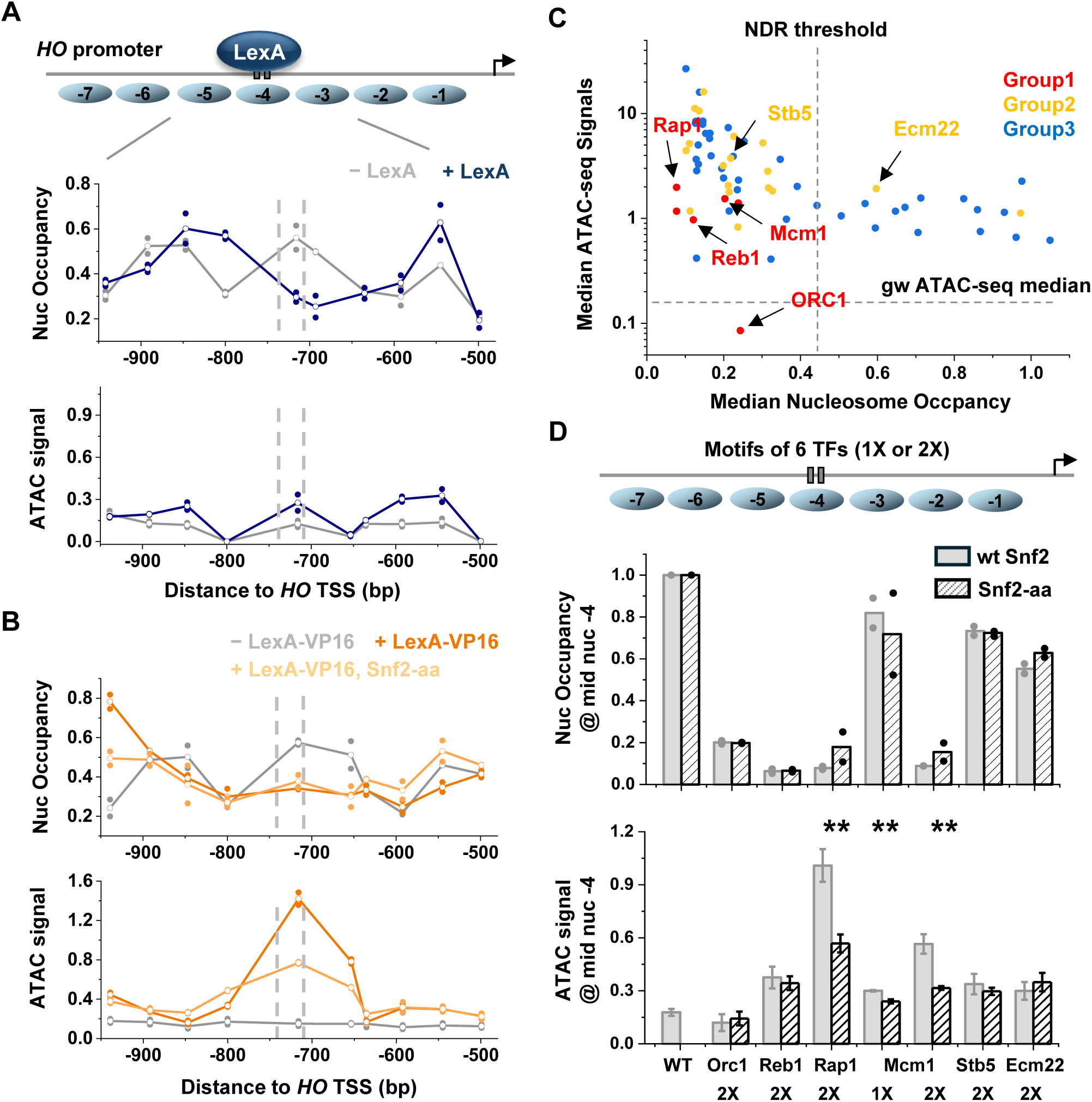
Transcription factors (TFs) differ in their ability to generate NDRs and ATAC-seq signal. **A)** LexA generates an ATAC− NDR. Top, schematic of a modified *HOpr* with two LexA binding sites engineered into nucleosome −4. Bottom, tiling MNase-qPCR and ATAC-qPCR across the nucleosome −5 to −3 region before (gray) and after (blue) LexA induction. The dashed line marks the position of the LexA binding sites. **B)** Same as in panel A, but with induction of LexA-VP16. LexA-VP16, but not LexA, strongly enhances ATAC-qPCR signal near the LexA binding sites. The light orange curve shows the same measurement after rapid Snf2 anchor-away following LexA-VP16 induction. **C)** Endogenous TF binding sites span a broad range of nucleosome occupancy and ATAC-seq signal. For each TF, the median nucleosome occupancy and median ATAC-seq signal across its genome-wide binding sites are plotted. **D)** Insertion of endogenous TF motifs into *HOpr* produces different effects on nucleosome occupancy and ATAC signal. For each factor, one motif (1X) or two motifs (2X) are inserted into nucleosome −4, and MNase-qPCR and ATAC-qPCR signals near the inserted motif are measured before and after Snf2 anchor-away.

Based on the results above that SWI/SNF promotes ATAC-seq signals, we next asked whether recruiting SWI/SNF is sufficient to convert a LexA-induced NDR into an ATAC+ NDR. We expressed LexA fused to VP16, a strong transactivation domain known to interact with SWI/SNF^41^. MNase measurements show that LexA-VP16 generates an NDR similar to LexA alone but also reduces the occupancy of the flanking nucleosomes (**Figure 6B**). Importantly, LexA-VP16 produces much stronger ATAC-qPCR signal near the NDR center (**Figure 6B**). Both the nucleosome occupancy and the ATAC signal partially revert back towards the LexA-only state when Snf2 is anchored-away (**Figure 6B**). Together, these results support a model in which VP16 recruits SWI/SNF to mobilize nearby nucleosomes and thereby generates strong ATAC signal.

We next examined endogenous yeast TFs. In a previous screen, we classified 104 sequence-specific TFs into groups 1, 2, and 3 because of their strong, weak, or undetectable nucleosome-displacing activity^42^. Because this classification is based entirely on MNase assays, it remains unclear how effectively these TFs can enhance ATAC-seq signal. To address this question, we analyzed MNase-seq and ATAC-seq profiles at TF binding sites. We focused on 67 factors for which published ChIP-exo data detected more than 10 binding events^17^. For each factor, we plotted the median nucleosome occupancy measured by MNase-seq against the median ATAC-seq signal across its genome-wide binding sites (**Figure 6C**), which reveals several distinct patterns. Orc1, which binds replication origins, is a clear outlier: its binding sites show low nucleosome occupancy, consistent with its classification as a group 1 factor^42^, but the corresponding ATAC-seq signal is below the background level. The remaining group 1 factors show low nucleosome occupancies and intermediate ATAC-seq signals, while group 2 and 3 factor binding sites show large variations in nucleosome occupancy, with medium to high ATAC-seq signals (**Figure 6C**). Thus, at TF binding sites, we can detect a partial disconnect between MNase-seq and ATAC-seq signals.

Because TFs often bind in clusters in the native genome^43^, the genome-wide analysis may not reflect the activity of individual TFs. To measure the chromatin-opening property of a single TF more accurately, we again used the synthetic system where we engineered motif(s) of a single TF into the *HOpr* and performed MNase and ATAC qPCR (**Figure 6D**). Based on the genome-wide pattern in **Figure 6C** and our previous TF studies^34,42^, we selected four group 1 factors, Orc1, Reb1, Rap1, and Mcm1, and two group 2 factors, Stb5 and Ecm22, for this test. For each of these factors, we inserted two motifs into *HOpr* nucleosome -4. We also engineered a single motif of Mcm1 10 bp from the nucleosome -4 dyad, a configuration previously shown to produce only weak nucleosome depletion.

Consistent with our previous MNase measurement, Mcm1 with a single motif, as well as the two group 2 factors, deplete nucleosome to much less extent in comparison with Orc1, Reb1, Rap1, and Mcm1 with two motifs (**Figure 6D & S6**). Importantly, the corresponding ATAC-seq signal is not simply the inverse of nucleosome occupancy. Orc1 generates an NDR but produces no ATAC-seq signal, consistent with the genome-wide pattern in **Figure 6C**. Stb5 and Ecm22 displace nucleosomes weakly but produce ATAC-seq signals comparable to that of Reb1. Rap1 and Mcm1 with two binding sites produce stronger ATAC-seq signals, which are partially dependent on SWI/SNF (**Figure 6D**). Together, these results indicate that native yeast TFs differ in their ability to generate NDRs and ATAC-seq peaks, which may be related to their abilities to recruit co-regulators to destabilize or mobilize the flanking nucleosomes.

### Discrepancies between ATAC-seq and MNase-seq are common among eukaryotes

We next examined previously published MNase-seq and ATAC-seq datasets from other species to determine whether the patterns observed in budding yeast are conserved across eukaryotes (**Resource Table**). We first analyzed the fission yeast *S. pombe*, another unicellular fungus that is evolutionarily distant from *S. cerevisiae*. We used the same pipeline applied to budding yeast to call NDRs and compare them with ATAC-seq peaks (**Methods**). This analysis again reveals a substantial discrepancy between ATAC-seq peaks and NDRs, identifying 788 overlapped regions, 2279 NDR only regions, and 202 ATAC only regions (**Figure 7A**). Consistent with budding yeast, most overlapped and NDR only regions are located in intergenic regions between tandem or divergent genes, while ∼60% of ATAC only peaks are located within gene bodies (**Figure 7B**). Overall, the differential pattern of ATAC-seq vs MNase-seq in fission yeast is highly similar to budding yeast.

**Figure 7.**
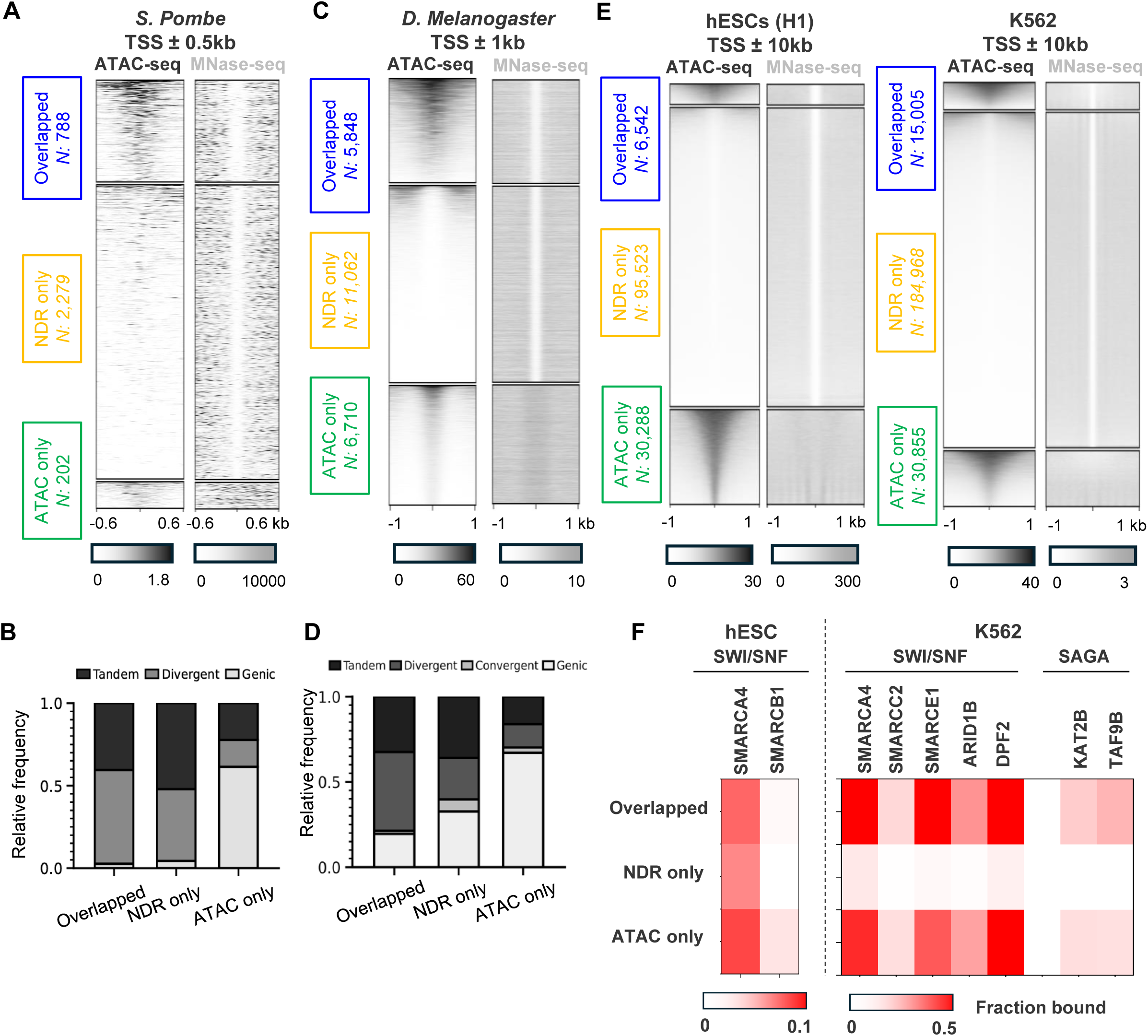
Discrepancies between ATAC-seq and MNase-seq are common among eukaryotes. **A)** Comparison and clustering of ATAC-seq peaks and NDRs in *S. pombe*. ATAC-seq peak calling, NDR annotation, and clustering are performed using the same approach as in Figure 1A. **B)** Genomic location of the overlapped, NDR only, and ATAC only regions in panel A. **C&D)** Same as in A & B, but in *D. Melanogaster*. **E)** Comparison and clustering of ATAC-seq peaks and NDRs in two human cell types, H1 embryonic stem cells and K562 cells. Both ATAC-seq peaks and NDRs are restricted to regions within 10 kb of annotated TSSs. **F)** Fraction of overlapped, NDR only, and ATAC only regions bound by SWI/SNF and SAGA subunits in hESCs and K562 cells.

We performed a similar analysis using ATAC-seq and MNase-seq in *Drosophila melanogaster* embryos collected at the 2-3 hrs after egg laying stage^44^. In *Drosophila*, nucleosomes in general adopt fuzzier positioning, but many promoters still contain well-defined NDRs upstream of TSSs^45^. We therefore restricted both ATAC-seq peak and NDR annotations to TSS ± 1kb. Intersecting ATAC-seq peaks with NDRs results in 5848 overlapped, 11062 NDR only, and 6710 ATAC only regions (**Figure 7C**). In comparison with the two fungi species, overlapped and NDR only regions in *Drosophila* are more likely to occur within gene bodies, but ATAC only regions still show the highest genic presence among the three groups (**Figure 7D**).

Finally, we examined ATAC-seq and MNase-seq datasets from human cells. We focused on H1 embryonic stem cells and the widely used K562 cancer cell line, both of which have high-quality ATAC-seq and MNase-seq data (**Resource Table**). We called NDR coordinates using the same method as in the other species, and restricted both NDRs and ATAC-seq peaks to regions within ±10 kb of annotated TSSs. In both cell lines, overlapped regions represent a minority of sites, with most regions classified as NDR only or ATAC only (**Figure 7E**). Overlapped regions are located closest to TSSs, with 86% and 70% falling within ±1 kb of annotated TSSs in H1 and K562 cells, respectively. In contrast, only ∼9% and 10% of NDR-only regions, and 64% and 47% of ATAC-only regions, are located within this promoter-proximal window in these two cell lines.

Given the strong enrichment of SWI/SNF and SAGA complex in yeast ATAC-seq peaks, we next asked whether similar enrichment is observed in human cells. SWI/SNF and SAGA ChIP-seq are not available for H1 cells, but SMARCA4 and SMARCB1 ChIP-seq data can be found in closely related pluripotent stem cell lines (H9 and iPSCs). ChIP-seq datasets for multiple SWI/SNF and SAGA subunits are available in K562 cells (**Resource Table**). Analysis of these datasets show that SWI/SNF and SAGA binding is generally higher in ATAC-seq peaks, including both overlapped and ATAC only regions, and this pattern is particularly prominent in K562 cells (**Figure 7F**). Together, these results indicate that SWI/SNF and SAGA enrichment at ATAC-seq peaks is conserved in human cells.

## Discussion

Chromatin accessibility is often treated as a single genomic property, yet our results show that MNase-seq and ATAC-seq capture distinct forms of accessible chromatin. Specifically, MNase-defined NDRs differ substantially from ATAC-seq peaks, which is observed across several distant eukaryotic species. The ATAC-seq pattern can be largely recapitulated by MNase-seq assay conducted at a lower concentration, indicating that intrinsic bias of MNase and Tn5 is only partly responsible for these differences. We found that the property of open chromatin plays an important role in this discrepancy: standard MNase-seq primarily identifies regions with stable nucleosome depletion, while ATAC-seq preferentially reports broader accessible chromatin that include dynamic nucleosomes. Thus, MNase-seq and ATAC-seq are not interchangeable readouts of a single open-chromatin state; instead, they emphasize different physical and functional properties of chromatin.

The comparison between high and low MNase digestion and ATAC-seq also provides insight into the concept of “fragile nucleosomes,” defined as regions that are protected from digestion at low MNase concentration but not at higher concentration. Because NDR only regions show discordant MNase-seq and ATAC-seq signals, one might expect them to contain partially protected fragile nucleosomes, whereas overlapped regions might represent fully open chromatin. Our data suggest the opposite: ATAC+ NDRs have higher MNase dose sensitivity than ATAC− NDRs (**Figure S3B**). Thus, by the operational definition of MNase-dose dependence, overlapped regions more closely resemble “fragile nucleosomes”, despite being strongly depleted of H3 (**Figure 1C**). In contrast, ATAC only regions, which harbor dynamic nucleosomes that can be considered as one type of “fragile nucleosomes”, show the lowest sensitivity to MNase concentration (**Figure S3B**). Together with the observation that overlapped regions bind more factors (**Figure 4B**), these results suggest that MNase-dose-dependent protection does not necessarily reflect fragile nucleosomes. Instead, it may primarily reflect partial protection by NDR-associated TFs and cofactors.

Although chromatin-associated factors are often assumed to bind within NDRs, our analysis identifies many factors that bind selectively inside ATAC-seq peaks, regardless of the extent of nucleosome depletion (**Figure 4B**). These ATAC-associated factors are enriched for transcriptional regulators and co-regulators, consistent with enhancer-associated activities. In contrast, factors enriched at NDRs include RNA polymerase II and general transcription factors, consistent with core promoter function. This distinction is notable because yeast lacks distal enhancers, and its upstream regulatory sequences tend to lie close to core promoters and are not clearly separated in gene annotations. Nevertheless, MNase-seq and ATAC-seq can distinguish these regulatory features. The ATAC+ vs ATAC-promoters are also connected to previously proposed “constitutive” versus “inducible” promoter classes (**Figure 4D**). Therefore, the difference between NDRs and ATAC-seq peaks reflects functional differences among open chromatin regions.

Importantly, the distinction between ATAC+ and ATAC− promoters should not be interpreted as a difference between active and inactive promoters. Promoters containing overlapped, NDR only, or ATAC only regions drive similar average transcription levels (**Figure 1F**). Although not directly tested here, prior literature on “constitutive” versus “inducible” promoters suggests that the key difference in transcription between these two classes is not absolute transcriptional output but in transcriptional plasticity. By recruiting co-regulators, ATAC+ promoters may support larger transcriptional changes under different growth conditions.

Through perturbation experiments, we also examined how several coactivators contribute to ATAC-seq signal. SWI/SNF promotes genome-wide ATAC-seq accessibility, whereas RSC has a smaller effect in the opposite direction. This result is surprising because SWI/SNF and RSC both belong to the SWI/SNF family of ATP-dependent chromatin remodelers, and both are described as nucleosome “pushers” that enlarge promoter NDRs^29^. Moreover, RSC is more abundant and essential, and its depletion causes widespread NDR shrinkage and nucleosomes to fill-in at more than 2,000 NDRs^29^. In contrast, SWI/SNF depletion alters the size of only ∼250 NDRs^29,34^. It is therefore often assumed that RSC acts broadly across the genome, whereas SWI/SNF functions more selectively through recruitment by sequence-specific TFs. Our ATAC-seq data challenge this view and suggest that nucleosome repositioning and Tn5 accessibility result from different remodeling activities. RSC seems to reposition nucleosomes while still protecting these nucleosomes from extensive Tn5 tagmentation. In contrast, SWI/SNF promotes Tn5 accessibility without inducing widespread nucleosome repositioning. The mechanism by which SWI/SNF destabilizes NDR-flanking nucleosomes remains to be determined, but possible models include increased DNA unwrapping, enhanced histone turnover, or transient nucleosome eviction. Overall, SWI/SNF and RSC appear to affect nucleosomes in qualitatively different ways, rather than simply acting at different genomic scales.

Our results also refine the concept of PF activity. Due to different usage of MNase-seq and ATAC-seq in different species, PFs in yeast are often defined as TFs that generates NDRs^42^, whereas pioneer activity in higher eukaryotes is more commonly evaluated by increased ATAC-seq signal after TF binding^46^. Using both artificial bacterial TFs and native yeast TFs, we show that TFs can have differential ability to generate these two forms of open chromatin. For example, LexA and Orc1 generate strong NDRs without increasing local ATAC-seq signal, whereas Stb5 and Ecm22 substantially increase ATAC-seq signal despite causing only weak nucleosome depletion. LexA-VP16, Rap1, and Mcm1 generate both NDRs and ATAC-seq signal, and this ATAC-associated activity is partially dependent on SWI/SNF.

Together with previous studies, these observations suggest that NDR formation and ATAC accessibility depend on distinct TF properties. LexA is not evolutionarily adapted for nucleosome binding and is unlikely to recruit native yeast chromatin regulators. Its ability to generate NDRs therefore suggests that strong DNA binding and high TF concentration can be sufficient to drive nucleosome depletion^42,47^. This behavior is consistent with a mass-action model in which TFs physically compete with histones for the same DNA sequence. It also agrees with our observation that endogenous yeast nucleosome displacing TFs tend to be abundant and have high DNA-binding affinity^42^. In contrast, strong ATAC-seq signals appear to depend more on recruitment of co-regulators that destabilize nearby nucleosomes and can arise even with weaker DNA binding / nucleosome depletion. Thus, TF regulatory domain can be particularly important for generating ATAC-seq peaks. This idea is consistent with previous reports that Grainy head (Grh) requires its disordered N terminus to induce ATAC-seq signal^48^, and that SWI/SNF inhibition causes a strong loss of GATA3-dependent ATAC-seq peaks^49^. Finally, motif configuration can also shape chromatin-opening activity: for some factors, such as Mcm1, single and multiple binding sites produce different levels of nucleosome depletion and ATAC-seq signal. Thus, depending on the assay and definition used, “pioneer activity” can reflect different TF properties and can also be highly site-specific.

## Methods

### Plasmid and yeast strains

Standard methods were used for strain construction and plasmid cloning. All yeast strains were derived from the W303 background. For TF-binding experiments, the endogenous *HO* promoter (*HOpr*; -1298 to -272 relative to the *HO* ORF) was deleted. A modified *HOpr* containing mutated Swi5-binding sites was then inserted into the *CLN2* locus. Consensus motifs for the TFs shown in **Figures 6 & S6** were engineered into nucleosome -4. The *MET3* promoter (*MET3pr*) was used to drive induction of LexA and LexA-VP16 in **Figures 6A & 6B**. Cells were grown overnight in SCD supplemented with 20X methionine. The following morning, cells were washed three times with water and resuspended in SCD lacking methionine to induce expression for 4 hours. After LexA and LexA-VP16 reached steady-state levels, Snf2 was depleted by anchor-away as described below.

For the Tra1 auxin-induced degradation strain, a 3X-V5-AID cassette containing 70-bp homology arms was PCR-amplified and integrated by homologous recombination to generate an in-frame gene fusion. Protein degradation was verified by treating cells with 500 µM 3-indoleacetic acid (Sigma-Aldrich, I2886) for 30, 60, or 90 minutes, followed by western blotting. The SWI/SNF and RSC anchor-away strains were generated using an FRB-TAP-GFP cassette to create in-frame gene fusions at the endogenous loci. Nuclear depletion was achieved by treating cells with 1 µg/mL rapamycin for 60 minutes and was confirmed by fluorescence microscopy. See **Table S1** for the complete primer and strain list.

### Western blotting

A total of 1.5 OD units of log-phase yeast culture were harvested, washed with 1 mL of ice-cold water, and resuspended in 200 µL of 1 M NaOH to lyse the cells. Samples were incubated at room temperature for 10 minutes. Cells were then pelleted, resuspended in 50 µL of SDS-polyacrylamide gel electrophoresis (SDS-PAGE) sample buffer, and boiled for 5 minutes prior to loading. To aid membrane transfer of Tra1-AID, transfer buffer was supplemented with 0.1% SDS and 10% methanol, and proteins were transferred overnight at 4 °C. Primary antibodies were used at the following dilutions: 1:5,000 anti-V5 (Abcam, ab27671) and 1:15,000 anti-α-tubulin (Abcam, ab184970). Secondary antibodies were used at the following dilutions: 1:3,000 goat anti-rabbit (Bio-Rad, 12004162) and 1:3,000 goat anti-mouse (Bio-Rad, 12004158).

### ATAC-seq

The ATAC-seq protocol used here was adapted from published Omni-ATAC protocol with slight modifications^50^. Briefly, yeast cultures were grown in SCD-met to an OD of 0.15, and 0.4 OD units were harvested per library. Cells were washed with 1 mL of 1 M sorbitol and incubated with 0.5 mg/mL zymolyase (E1005) for 7 minutes at room temperature with gentle inversion. Cells were then washed with 1 mL of 1 M sorbitol and incubated with 2 µL of Diagenode Tn5 (C01070013-80) in 48 µL of 1× TD buffer (25 µL of 2× tagmentation buffer (C01019043-1000) supplemented with 0.01% digitonin, 0.1% NP-40, and 0.1% Tween-20) at 37 °C for 30 minutes with mixing on a nutator. Samples were purified using the E.Z.N.A. Cycle-Pure purification kit (101318-890). PCR was performed using NEBNext High-Fidelity 2× PCR Master Mix (M0541S). To avoid overamplification, sequencing libraries were amplified using a two-step process as described previously^3^. Briefly, an initial 50-µL PCR reaction was performed for five cycles. Samples were then placed on ice, and 5 µL was used in a qPCR reaction to determine the number of additional cycles required to reach one-third of the PCR plateau (typically seven additional cycles). Once the cycle number was determined, the remainder of each sample was amplified accordingly, purified using NEB sample beads at a 1.2× bead-to-sample ratio, and quantified by TapeStation. For ATAC-qPCR, tiling PCR was performed over the *HOpr* (**Table S1**) using the amplified sequencing libraries. Primers flanking the TF binding site(s) were used to quantify enrichment relative to a genomic control site with strong ATAC signal. ATAC-qPCR signal was calculated as fold-change relative to an un-tagmented genomic sample (input) and then normalized to the positive control (**Table S1**). For ATAC-seq, all libraries were sequenced on an Illumina NextSeq 2000 using 2 × 50-bp paired-end reads.

### MNase-seq

MNase assay was performed similar to previously described^42^. Briefly, 10ml cells with 1.5 OD660 0.15 were harvested and washed sequentially with 1 mL of water and 1 mL of 1 M sorbitol. The pellet was resuspended in 0.5 mL of spheroplasting solution (1 M sorbitol, 0.5 mM 2-mercaptoethanol, and 0.18 mg/mL zymolyase) and incubated at room temperature for 7 minutes with gentle inversion. Spheroplasts were collected by centrifugation at 500 × g for 3 minutes, washed twice with 1 mL of 1 M sorbitol, and resuspended in 200 µL of digestion buffer (1 M sorbitol, 50 mM NaCl, 100 mM Tris-Cl pH 7.4, 5 mM MgCl2, 1 mM CaCl2, 1 mM 2-mercaptoethanol, 0.5 mM spermidine, and 0.075% NP-40). Samples were incubated at 37 °C for 8 minutes with 1 U of MNase (Sigma N3755). Reactions were terminated by adding 20 µL of quench buffer (250 mM EDTA and 5% SDS). DNA was purified by phenol-chloroform extraction, and mononucleosome-sized DNA was gel-extracted before proceeding to stacking qPCR. Nucleosome occupancy was normalized to a reference nucleosome in the EXO84 terminator.

MNase-seq follows the same procedure, with 1.5 OD660 units of cells were digested with 0.1 U or 1 U of MNase (Sigma N3755) for 8 minutes, resulting in approximately 30% or 70% of the genome being converted into mononucleosomes, respectively. A total of 100 ng of MNase-digested DNA was used to generate sequencing libraries with the NEBNext® Ultra™ II DNA Library Prep Kit for Illumina® (Catalog # E7645L), following the manufacturer’s protocol. Libraries were sequenced on an Illumina NextSeq 2000 in 2 × 50-bp paired-end mode.

### ATAC-seq data analysis

∼10 million reads were generated for each ATAC-seq experiment. Reads were aligned to the sacCer3 genome using Bowtie 2 with the “—very-sensitive” preset. Alignment files were filtered using SAMtools to remove unmapped and unpaired reads, as well as alignments with MAPQ scores below 30. PCR duplicates were identified and removed using samtools markdup, and reads aligned to repetitive regions or the mitochondrial genome were blacklisted and excluded. ATAC-seq peaks were called using MACS2 (version 2.2.9.1) in paired-end mode. Peak calling was performed using a q-value of e^-4 across the three individual replicates and for the merged replicates and consensus peaks were then selected as regions that were called across the three replica (*N* = 1627). Peaks were filtered to remove peaks overlapping any blacklisted regions or intergenic convergent regions. Finally, peaks were further filtered by requiring each peak to be within 500 bp of a TSS (*N* = 1242). Genome browser visualizations were performed using IGV. When ATAC-seq datasets were directly compared, BigWig files were normalized to 1× genome coverage using deepTools. Sites of Tn5 insertions were identified by shifting the left and right ends of each aligned read inward by 4bp to account for Tn5 binding^3^. Fragment sizes were determined from filtered bam files using the absolute paired-end template length (TLEN).

### MNase-seq data analysis

∼20 million reads per sample were generated and aligned to the sacCer3 genome assembly using Bowtie2 with the “--very-sensitive” preset. BAM files were filtered to remove unmapped and unpaired reads, as well as alignments with MAPQ scores below 30. Biological replicates were merged prior to downstream analysis. BigWig files were generated using deepTools bamCoverage (version 3.5.6), with fragments size from 100-200 bp unless otherwise stated. Alignments were trimmed by 15 bp from each end to sharpen the nucleosome occupancy. NDRs were identified using a genome-wide MNase signal threshold. By default, the 12.5th percentile of genome-wide MNase signal was used to define low-coverage positions. Each chromosome was scanned using a 75-bp sliding window, and a window was classified as nucleosome-depleted if no more than 20 bp exceeded the genome-wide signal threshold. Nucleosome-depleted windows separated by 105 bp or less were merged into a single NDR. Regions greater than 1,000 bp prior to merging were excluded. NDRs overlapping blacklisted regions or the mitochondrial genome were excluded. Nucleosome positions and fuzziness scores were calculated using DANPOS3^22^ with default parameters.

### ATAC and MNase comparisons

Because budding yeast terminators generally exhibit weak nucleosome depletion, NDRs and ATAC-seq peaks located more than 500 bp from an annotated TSS or within convergent intergenic regions, were excluded from subsequent analyses. The remaining regions were intersected using BEDTools intersect, generating three clusters in **Figure 1A**. To facilitate direct comparison between overlapping and NDR only regions, overlapping regions were recentered using the coordinates of the corresponding NDR for downstream analyses and visualization. Genes were identified using RefSeq annotations, and the genomic coordinates and strand orientation of neighboring genes were used to classify tandem vs divergent genes. Heatmaps, including those shown in **Figure 1A & C**, were generated using deepTools. The nucleotide composition of each region was determined from the *S. cerevisiae* reference genome using BEDTools nuc.

### Perturbation analysis

Average ATAC-seq change in **Figure 5C** was quantified based on the sequencing coverage within 100 bp windows centered at the midpoints of NDRs in the overlapped and NDR only regions, and midpoints of ATAC-seq peaks in the ATAC only regions. ATAC-seq coverage was then averaged across all regions within each cluster. Normalized ATAC-seq signal in the overlapped and ATAC only regions were calculated as fold change relative to that of the NDR only regions, and compared with a WT sample.

### Yeast promoter classification

Promoter classifications with corresponding NDR and TSS coordinates were downloaded from (PMID: 36302553) Supplemental Table S1. Promoters were classified as ATAC+ or ATAC-based on whether they contain ATAC-seq peaks. Overlap between ATAC+ or ATAC-promoters with additional genomic features defined in **Figure 4D** were determined by BEDTools intersect.

### Factor binding analysis

To determine enriched TF binding in the different regions, we downloaded the complete set of ChIP-exo binding sites from the Yeast Epigenome Project^17^. For analysis in **Figure 4**, we calculated the fraction of binding events for each factor located in overlapped, NDR only, and ATAC only regions (center ± 100bp). Factors were then clustered using the kmeans function in MATLAB, with the optimal number of clusters determined by the elbow method. For analysis in **Figure 5A**, we performed the same binding-probability calculation using AT-content- and length-matched “overlapped” and “NDR only” regions. We then calculated the log2 fold enrichment between the two region classes and assessed significance using a two-proportion Z-test.

### TSS analysis

Transcription start site data for *S. cerevisiae* was obtained from the YeasTSS database for log phase cells grown in YPD^28^. Overlapped, NDR only, and ATAC only sets were oriented relative to their closest TSS and used for plotting the TSSs distribution in **Figure 3D**.

### S. pombe analysis

ATAC-seq data were obtained from GSE66386^13^ and aligned to the *S.pombe* ASM294v2 reference genome using Bowtie2 v2.5.4 with the --very-sensitive preset. Alignments were filtered using SAMtools v1.22.1 to retain only properly paired reads (-f 2) with a mapping quality ≥30 (-q 30), while excluding reads mapping to mitochondrial reads. PCR duplicates were identified and removed using samtools markdup. Following filtering and deduplication, biological replicates were merged and ATAC-seq peaks were called using MACS2 in paired-end mode with a q-value threshold of e^-4 (*N* = 1721). Genome-wide ATAC-seq coverage tracks were generated from fragments 30–130 bp in length using bamCoverage from deepTools v3.5.6. MNase-seq data were obtained from GSE227182^51^, and genome-wide NDRs were called using the same algorithm as in budding yeast (*N* = 5230). We then selected ATAC-seq peaks and NDRs that are located from −500 to +500 bp relative to TSSs^52^ (*N* = 914 & 3118). These TSS-proximal ATAC-seq peaks and NDRs are intersected to generate **Figure 7A**.

### *Drosophila* embryo analysis

MNase-seq and ATAC-seq datasets from *Drosophila melanogaster* were obtained from Brennan et al.^44^. For MNase-seq, WT embryo samples corresponding to 2–3 h after egg laying (SRR5486181 and SRR5486184) were used. For ATAC-seq, untreated embryo samples from the corresponding developmental interval (SRR7813063–SRR7813066) were used. Reads were aligned to the *D. melanogaster* dm6 reference genome using Bowtie2 with the --very-sensitive settings. Alignments were filtered using SAMtools v1.22.1 to retain only properly paired reads (-f 2) with a mapping quality greater than 30. Mitochondrial reads were excluded, and PCR duplicates were identified and removed using samtools markdup. Peaks were then called with MACS2 in paired-end mode using a q-value threshold of e^-4 (*N* = 28,837). ATAC-seq peaks were then filtered to remove blacklisted region and selected for those that were within 1 kb of a TSSs (*N* =11,608). Genome-wide NDRs were called using the same program used above (*N* = 59,078) and NDRs were retained for those that were within 1 kb of a TSSs (*N* = 16,910). ATAC peaks and NDRs were then intersected to generate **Figure 7C**.

### Human cell analysis

H1 hESCs and K562 analyses were based on the hg19 genome assembly. All the data used are listed in the Resource Table. ATAC-seq peaks are called in MACS2 using standard settings. ATAC-seq data sets for H1and K562 were aligned to hg19 using Bowtie2 with the –very-sensitive settings. The alignment file was sorted and indexed and reads with MAPQ scores less than 30 and mitochondrial reads were removed. Duplicates were marked and removed with samtools markdup. DeepTools alignmentSieve –ATACshift was used to generate cut-site Bam files. Bam files were then merged with samtools merge and bigwig files generated with deepTools bamcoverage. NDRs were identified using the same algorithm as above. Both ATAC-seq peaks and NDRs were further filtered to remove any blacklisted regions and retain only those within 10 kb of an annotated TSS (UCSC hg19 refGene) resulting in 102,065 NDRs and 36,830 ATAC peaks for H1 hESCs and 199,973 NDRs and 60,228 ATAC peaks in K562. **Figure 7D** is generated by intersecting the filtered ATAC-seq peaks with filtered NDRs.

## Data availability

ATAC-seq datasets have been deposited at GEO under the accession GSE346229 and MNase-seq datasets have been deposited at GEO under the accession GSE346150. Previously published datasets used in this paper have been listed in the resource table.

## Acknowledgements

We thank Dr. David Stillman for providing the nhp6a/b double deletion strain. We acknowledge the Huck Institutes’ Genomics Research Incubator (RRID:SCR_023645) for use of the NextSeq 2000 and Cheryl Keller for help with sample preparation. We acknowledge all members in the Bai lab for insightful comments on the manuscript. We also thank the members of the Center of Eukaryotic Gene Regulation at Pennsylvania State University for discussions. This work is supported by the National Institutes of Health (T32 GM125592 to M.G. and R35 GM139654 to L.B.)

