## Supplementary_figure for "ATAC-seq and MNase-seq Detect Distinct Modes of Chromatin Accessibility"

### Figure S1

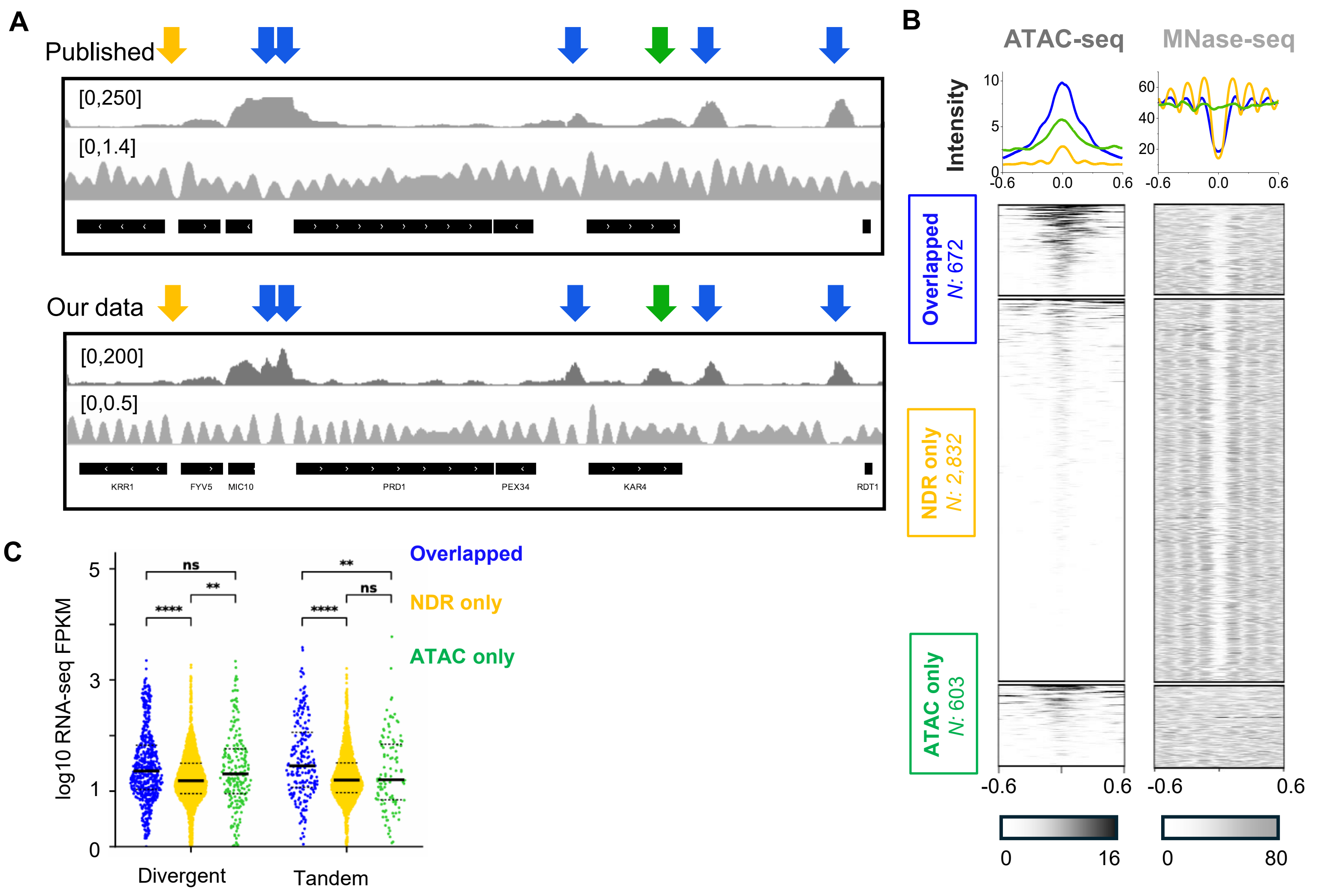


**Figure S1. More data related to ATAC-seq peak and NDR companiosn. A)** Additonal example of overlapped (blue), NDR-only (orange), and ATAC-only (green) regions. The top and bottom panels show the same genomic loci using published data[13,14] and our own data, respectively. **B)** Heat map of published dataset[13,14] using the same region coordinates in Figure 1A. **C)** Same analysis as in Figure 1F, performed using RNA-seq (steady-state RNA)[21] instead of RATE-seq (nascent RNA) data. ns: not significant, *P*-value < 0.01 (*), < 0.001 (**), <0.00001 (****).

**Figure S2**

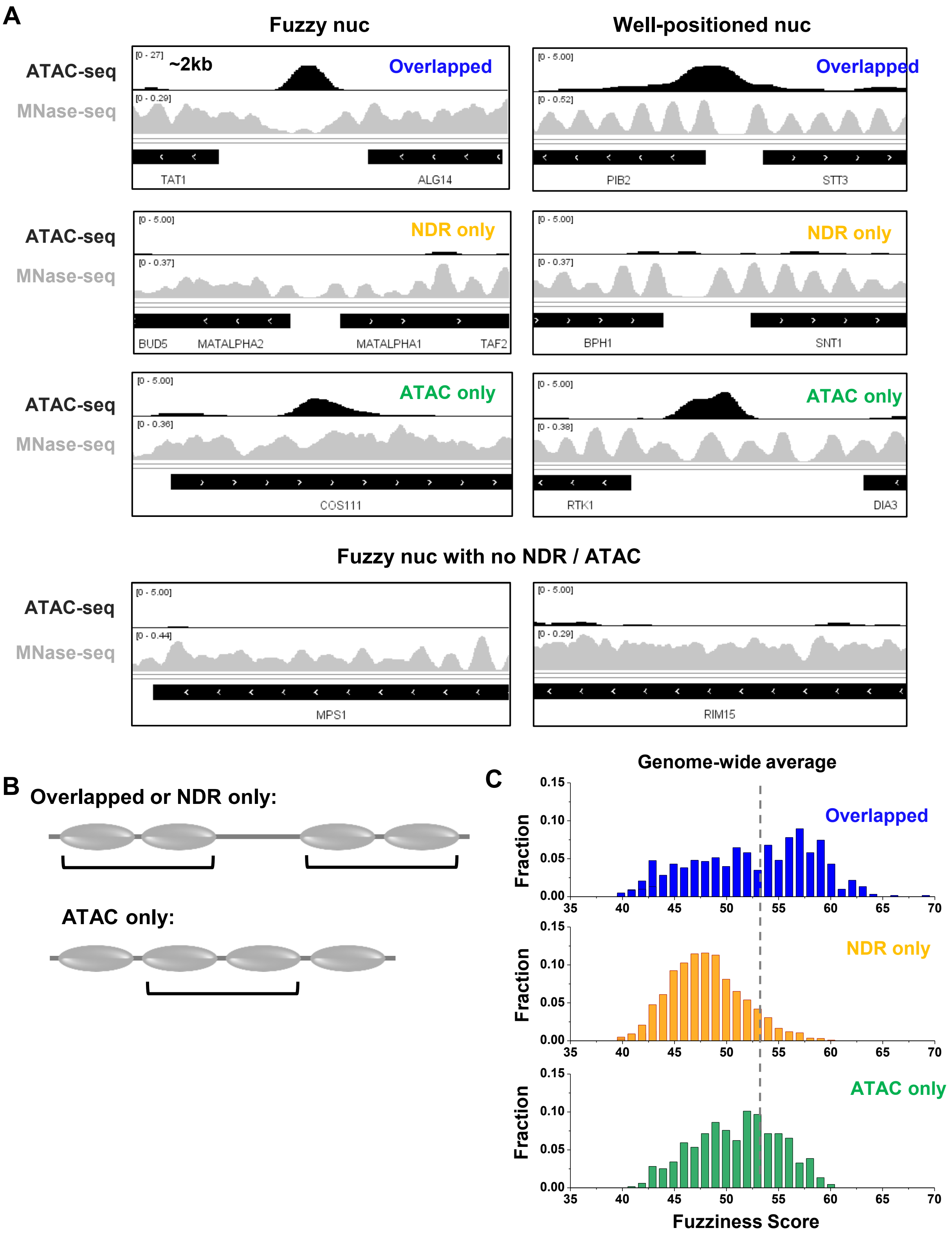


**Figure S2. ATAC signals are not caused by "fuzzy" nucleosome positioning. A)** Top three rows, examples of overlapped, NDR-only, and ATAC-only regions located near fuzzy (left) or well-positioned (right) nucleosomes. Bottom, examples of fuzzy nucleosome arrays that lack both ATAC-seq signal and NDRs. **B & C)** DANPOS analysis[22] of nucleosome fuzziness score. For overlapped and NDR-only regions, fuzziness scores were calculated for the four nucleosomes flanking each NDR, two on each side. For ATAC-only regions, fuzziness scores were calculated for the two nucleosomes at the center of each ATAC-seq peak. The distribution of fuzziness scores is shown in panel C.

Figure S3

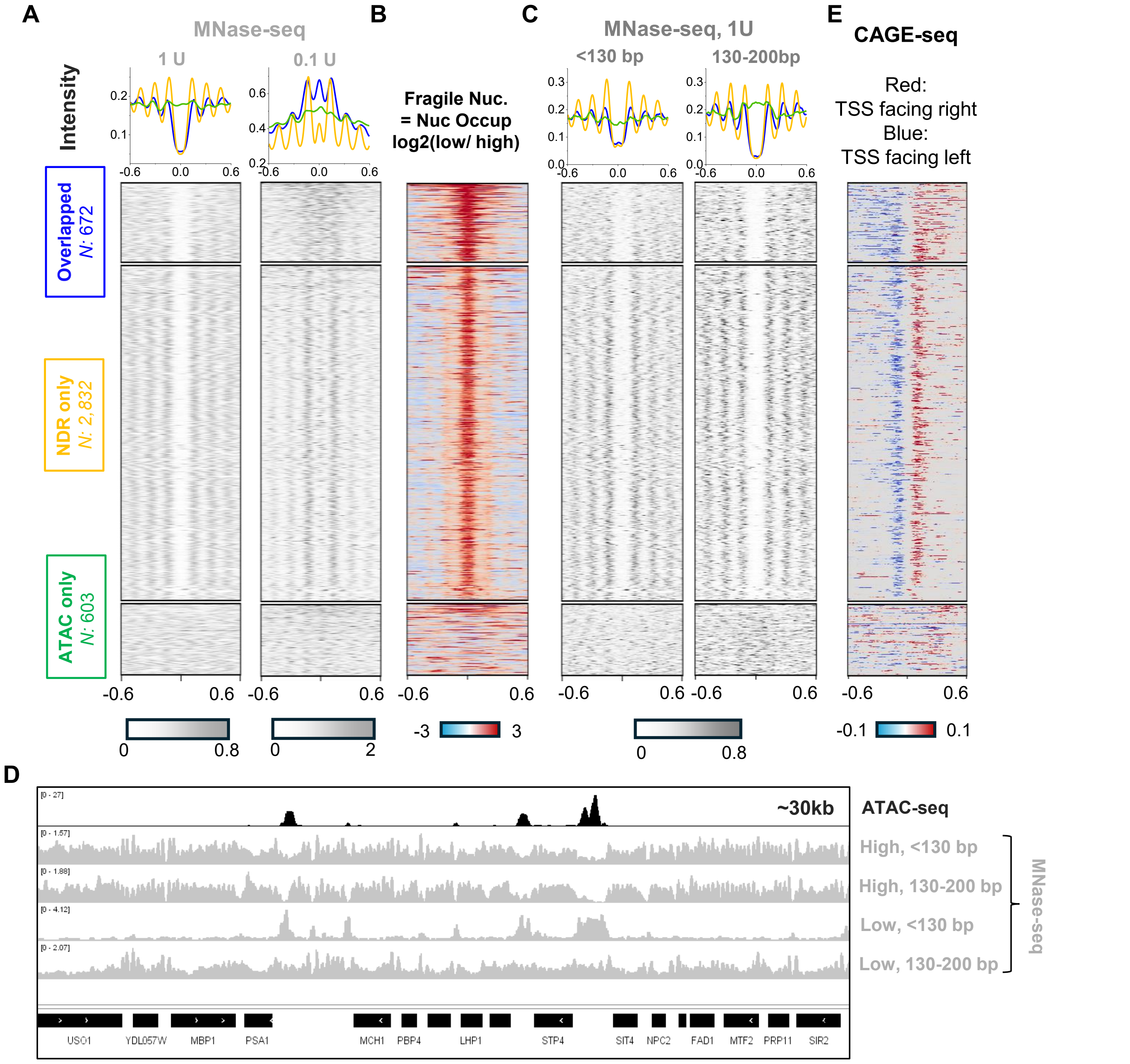


**Figure S3. Additional MNase analysis. A)** Aggregate MNase-seq profiles generated without fragment-size selection under high- and low-digestion conditions, using 1 U and 0.1 U MNase, respectively. **B)** Log2 fold difference between low- and high-MNase-seq signals. Higher values here are often used as an indicator of fragile nucleosomes. **C)** High-MNase-seq data separated by fragment size. Left: sub-nucleosomal (<130 bp), right: nucleosmsal (130-200 bp). **D)** Representitive genomic tracks of ATAC-seq and size-selected MNase-seq data generated under high- and low-MNase conditions. Note the similarity between ATAC-seq signal and the subnucleosomal fraction in the low-MNase data. **E)** CAGE-seq data[28] across overlapped, NDR-only, and ATAC-only regions.

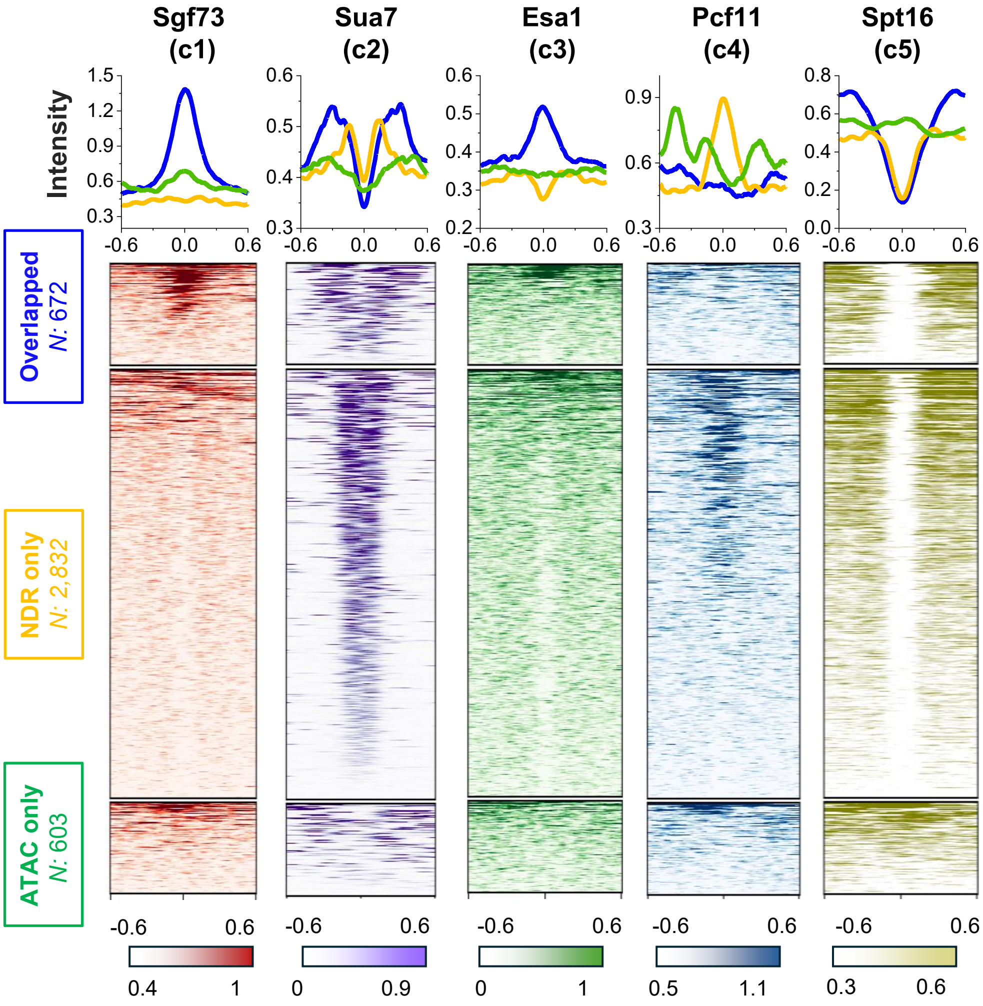


**Figure S4. Examples of chromatin-associated factors from clusters c1–c5.** The clusters of different binding patterns are defined in Figure 4B.

### Figure S5

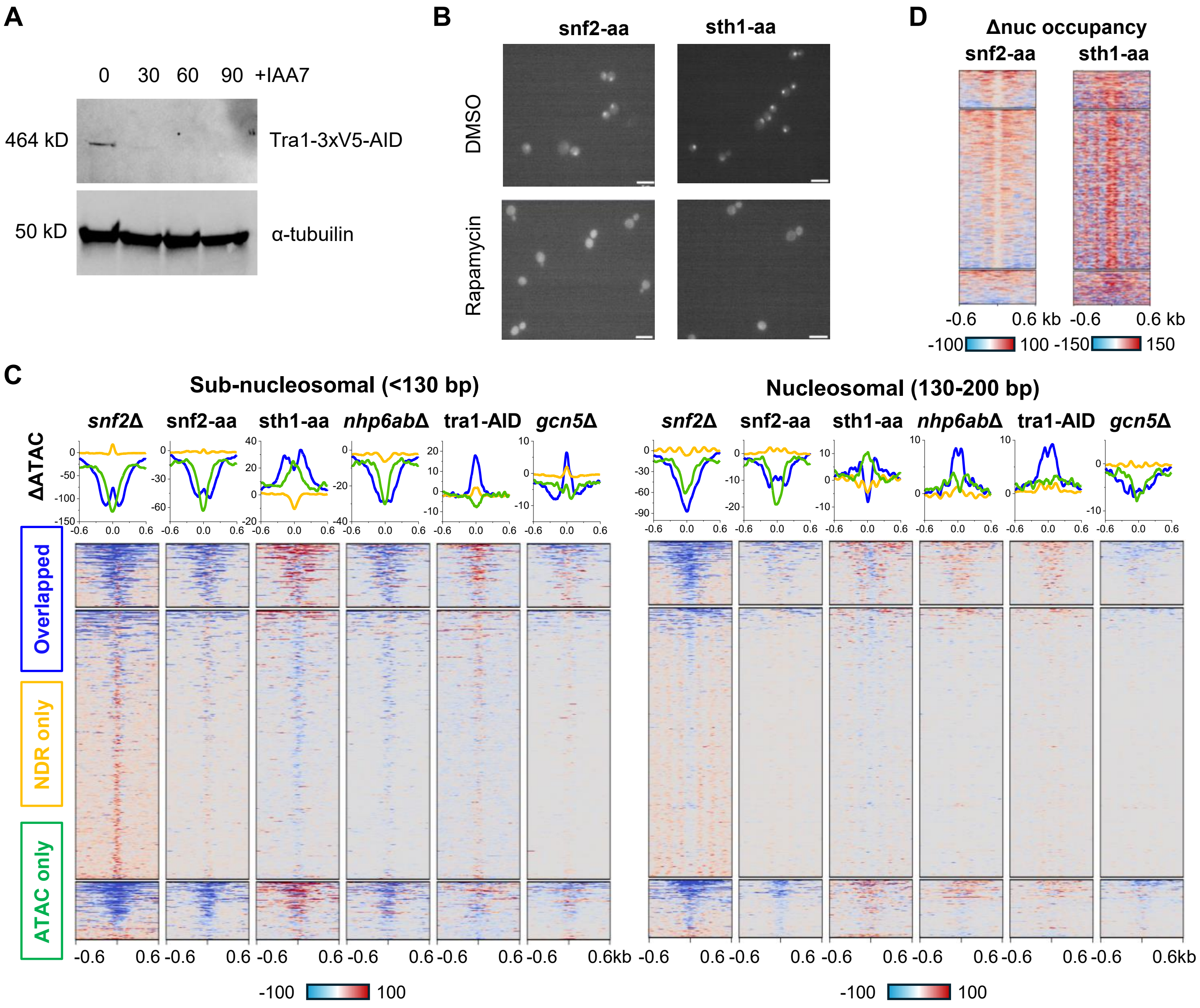


**Figure S5. Supplementary data for perturbation experiment in Figure 5. A)** Western blot confirming the auxin-induced degradation of Tra1. **B)** Imaging analysis confirming anchor-away depletion of Snf2 and Sth1. Both factors lose nuclear localization after rapamycin treatment. Scale bars are 10 μm. **C)** Changes in ATAC-seq signal following disruption of selected factors. The same data as in Figure 5B are shown, except that ATAC-seq reads here are separated by fragment size. **D)** Changes in nucleosome occupancy following Snf2 or Sth1 anchor-away. The Sth1 data are from our lab[40], and the Snf2 data are from literature[29].

**Figure S6**

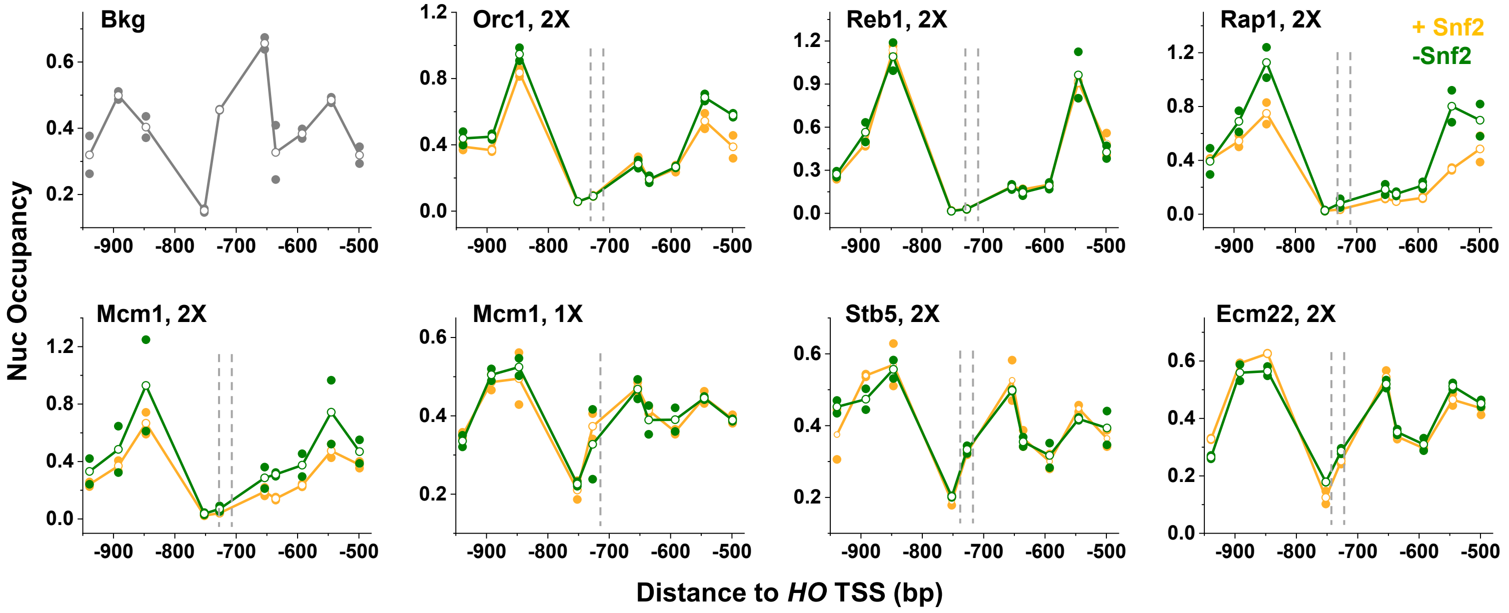


**Figure S6. Tiling MNase-qPCR over *HOpr* containing different TF motifs.** TF identities and motif copy numbers are indicated in each panel. The two curves show measurements before (orange) and after Snf2 anchor-away (green).
